# Comparative Transcriptomics of heads and tails between *Steinernema carpocapsae* and *Caenorhabditis elegans*

**DOI:** 10.1101/2020.08.20.259739

**Authors:** Isaryhia Maya Rodriguez, Lorrayne Serra Clague, Cassandra Joan McGill, Bryan Rodriguez, Ali Mortazavi

**Author notes:** These Authors contributed equally to this work (Isaryhia Maya Rodriguez, Lorrayne Serra Clague).

## Abstract

*Steinernema nematodes* have been widely studied for insect infection and mutualism, but little is known about the patterns of gene expression along the body of these worms or how these compare to the model organism *Caenorhabditis elegans*. Here we perform the first comparative analysis between the heads and tail regions of *Steinernema carpocapsae* and *C. elegans* Infective Juveniles (IJs)/dauers and young adults using single-worm RNA-seq. While we find overall agreement in gene expression there were several sets of genes with substantial differences between the two species. Gene expression in the *S. carpocapsae* female compared to the *C. elegans* hermaphrodite heads and tails revealed differences in metabolism, aging, and determination of lifespan. Young adult male heads and tails showed major differences in developmental related processes such as morphogenesis as well as neuronal development and signaling. We also found head- and tail-specific gene expression differences between *S. carpocapsae* IJs and *C. elegans* dauers for genes related to growth and development as well as neuronal signaling and activity. This study is one of the first comparative transcriptomic analyses of body parts between distantly related species of nematodes and provides insight into both the highly conserved and genetically distinctive characteristics of both species.

## Introduction

Nematodes are a diverse group of metazoans divided into 12 clades, including both free-living and parasitic species, which inhabit nearly every known biological niche on the planet^1,2^. Nematodes have been studied for their diversity, their roles as meiofauna in many ecosystems, the evolution of parasitism within the phylum, and parasitic infections in plants, animals, and humans alike^3–6^. The study of nematodes has provided insights into fundamental human genetic and molecular functions, potential application as biocontrol, and agricultural impact^7–9^. Despite their extremely high level of diversity and abundance on the planet, while some nematode clades are exceptionally well characterized through individual and comparative study, others have been significantly less explored and have a high potential for insight into nematode evolution, parasitism, and development as a complementary biological model.

One nematode genus, which is primarily studied for its biological control capabilities, is *Steinernema*. The Steinernematidae family consists of Entomopathogenic Nematodes (EPNs) that efficiently infect and kill insects. EPNs are obligate insect parasites, and for that reason, they have been used as biocontrol for many insect pests^10^. EPNs can only infect and kill an insect during their infective Juvenile (IJ) stage. IJs are characterized by a stop in development where the nematode can survive outside of a host for months and is equivalent to the dauer stage in *C. elegans*. EPNs such as *Steinernema feltiae* and *S. carpocapsae* can be bought commercially as organic alternatives to pesticides and are known as beneficial nematodes^11^. In addition to biocontrol, EPNs have also emerged as a model organism for mammalian parasites^12^. While human pathogens are more dangerous and challenging to maintain in a lab setting, EPNs are an advantageous alternative from which we can safely learn more about parasitic mechanisms and behaviors.

Within the Steinernematidae family, *S. carpocapsae* serves primarily as a beneficial nematode but has the potential to reveal conserved mechanisms across EPNs and mammalian parasitic nematodes^12^. *S. carpocapsae* is gonochoristic, having distinct males and females (Fig. 1A). The adult male has a tail with an evident spicule for mating and gonads that follow the same formation pattern of *C. elegans* males^13^. On the other hand, the adult females are larger than the males by at least 3 times and have variability in gonad arm migration^13,14^. While less is known about female and male adults, IJs have been extensively studied and provided insightful information into EPNs’ venom production and its mutualistic bacteria, *Xaenorhabditis nematophila*, that is dispensable for infections^15–17^. Transcriptome comparison between *S. carpocapsae* and *S. feltiae* exposed a shared set of core venom proteins and led to a study of highly conserved FMRFamide-like peptides, found in *C. elegans*, that was revealed to be essential for sensory perception in *S. carpocapsae*^12,18^. *S. carpocapsae* adult females, males, and IJs offer important insight into parasite development, including anatomical structures that are critical for parasitism, development inside the host, and mating.

**Figure 1.**
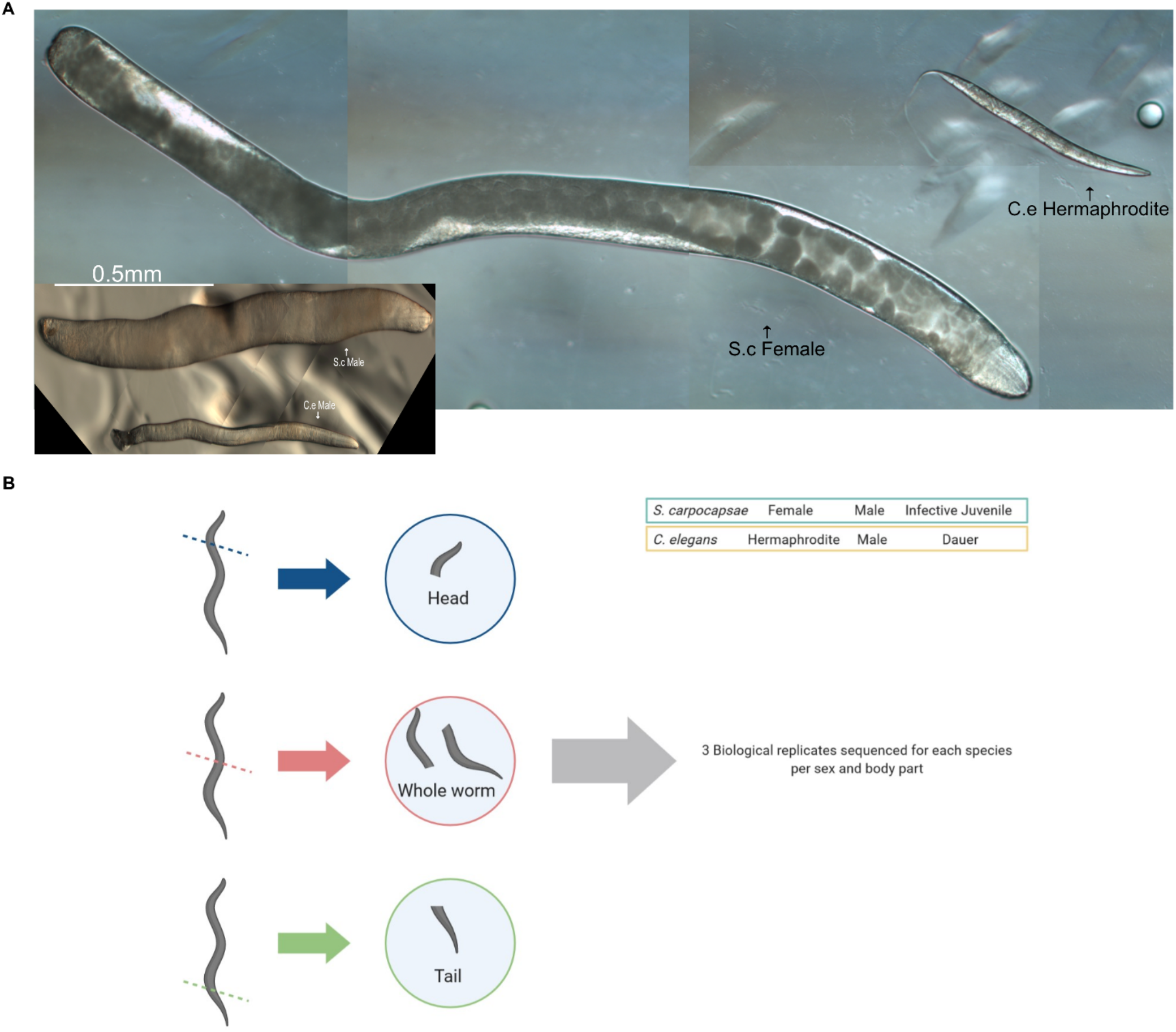
Comparison of size and morphological structures between *S. carpocapsae* and *C. elegans* equivalent sex and basic experimental design. (a) (Right) *S. carpocapsae* adult female in comparison to *C. elegans* adult hermaphrodite in bright field (50x). To the left of S.c. female is *S. carpocapsae* adult male in comparison to *C. elegans* adult male (200X). Photos have been scaled to demonstrate comparative size across sexes. (b) Cartoon of basic experimental design detailing cuts on worms and three biological replicates of each type of sample collected including head (blue), whole worm (pink) and tail (green) which were then isolated and lysed to extract mRNA using polyA selection and sequenced on the Illumina platform. Created with BioRender.com.

*C. elegans* is an excellent model organism for comparison, useful for obtaining insights into *S. carpocapsae* IJ and adult transcriptomes. The *C. elegans* genome, interactomes, and transcriptomes paved the way for comparative analysis with closely related species such as *Caenorhabditis briggsae* and more distantly related species such as *Pristionchus pacificus* and *Toxocara canis*^19–22^. Comparisons between closely related species offered insights into genetic variation in selfing and population genetics, while comparative analyses between distantly related species revealed conserved developmental and transcriptional programs for *C. elegans* dauer related genes in *P. pacificus*^21^. Additionally, the same dauer related genes in *C. elegans* are essential for development and mammalian host migration in *T. canis*^22^. Ultimately, comparative analysis of *C. elegans* to other species has provided valuable information for population genetics, gonad development, genetic variation, and conserved developmental processes across multiple nematodes’ clades and genera.

Here, we report a comparative transcriptome analysis between *C. elegans* young adult male, young adult hermaphrodite, and dauer to *S. carpocapsae* young adult male, young adult female, and IJ stages. To further understanding of anatomical structures in development, we analyzed the head and tail, as well as the whole animal. To perform the transcriptome analysis, we used single nematode RNA-sequencing and obtained high-resolution data of gene expression^23^(Fig. 1B). We first do comparisons within species before comparing across species. This study is the first to analyze region-specific transcriptomic differences across distantly related nematode species.

## Results

### *C. elegans* region-specific transcriptomic comparison of young males and hermaphrodites

Region-specific transcriptomic comparative analyses have been performed extensively in *C. elegans*^24–26^, and thus offers a valuable platform to further our understanding of *S. carpocapsae* young adult and IJ stages. In order to compare the region-specific transcriptomes between *C. elegans* and *S. carpocapsae*, we tested first whether our single-nematode RNA-seq method can detect known, region-specific expression in *C. elegans*. We therefore collected triplicates of *C. elegans* whole worms, heads, and tails for two stages: dauers and young adult (both hermaphrodites, and males). We detected an average of 18,414 genes in whole young adult hermaphrodites (at least one mapped read) and 19,425 genes in whole young adult males, which is equivalent to 91% and 96%, respectively, of genes identified in bulk nematode sequencing previously done in the same stage^27^. We measured average TPM values in hermaphrodite heads and male heads for the gene *myo-2*, a protein crucial for actin filament binding activity, and *flp-1*, an FMRF-like peptide, which is expressed in the nerve ring^28,29^. As expected, *myo-2* and *flip-1* are expressed in the whole young adult hermaphrodite and male, and in both heads but not in the tails (Fig. S1A, B). For hermaphrodite tail-specific expression, *egl-20*, a tyrosine kinase part of the Wnt signaling pathway, is found in the whole young hermaphrodite and tail but not in the head, as expected^30^ (Fig. S2A). Lastly, polycistine-1 (*cwp-1*), expressed by male-specific B type sensory neurons in the head and tail, *ins-31*, insulin related and expressed in male gonad, and *clec-207*, C-type lectin present in the vas deferens, are all found in the tail region of the male, except for *cwp-1* which as predicted is also expressed in the head of males^27,31,32^ (Fig. S2B). As anticipated, the expression of these genes corresponds to their anatomical localization, therefore validating our method of collection.

We then performed a comparative analysis between *C. elegans* young adult hermaphrodite and a young adult male. We found 8,334 genes that were characterized as differentially expressed (DE) in at least one tissue between young adults, and young adults versus dauers (counts per million (CPM) ≥ 2, FDR ≤ 0.05 and logFC ≥ 2). Whole young adults, heads, and tails have distinct transcriptome profiles, sharing only a small subgroup of genes (Fig. 2A). The comparative gene expression analysis of *C. elegans* tissues revealed 2,217 genes enriched in whole young adult hermaphrodites and 877 genes in whole young adult males (Fig. 2B, Table S1). The biological process gene ontology (GO) analysis for whole young adult hermaphrodites indicates terms involved in morphogenesis (*post-embryonic body morphogenesis, developmental processes, molting cycle*), and body and cell wall composition (*cell wall organization, extracellular matrix, aminoglycan catabolic activity*). Meanwhile, GO terms for whole young males were enriched for neural related terms (*neuropeptide signaling pathway, chemical synaptic transmission, potassium ion transport, cell projection*), reproduction (*gamete generation, cilium organization, phosphorus metabolic process*) and catabolic process (*cellular, nitrogen compound, cellular aromatic compound*) (Fig. S2, Table S2). The GO analysis for both sexes is related to young adult tissue that is in the process of finishing the development of reproductive structures, which agrees with recent findings^33^.

**Figure 2.**
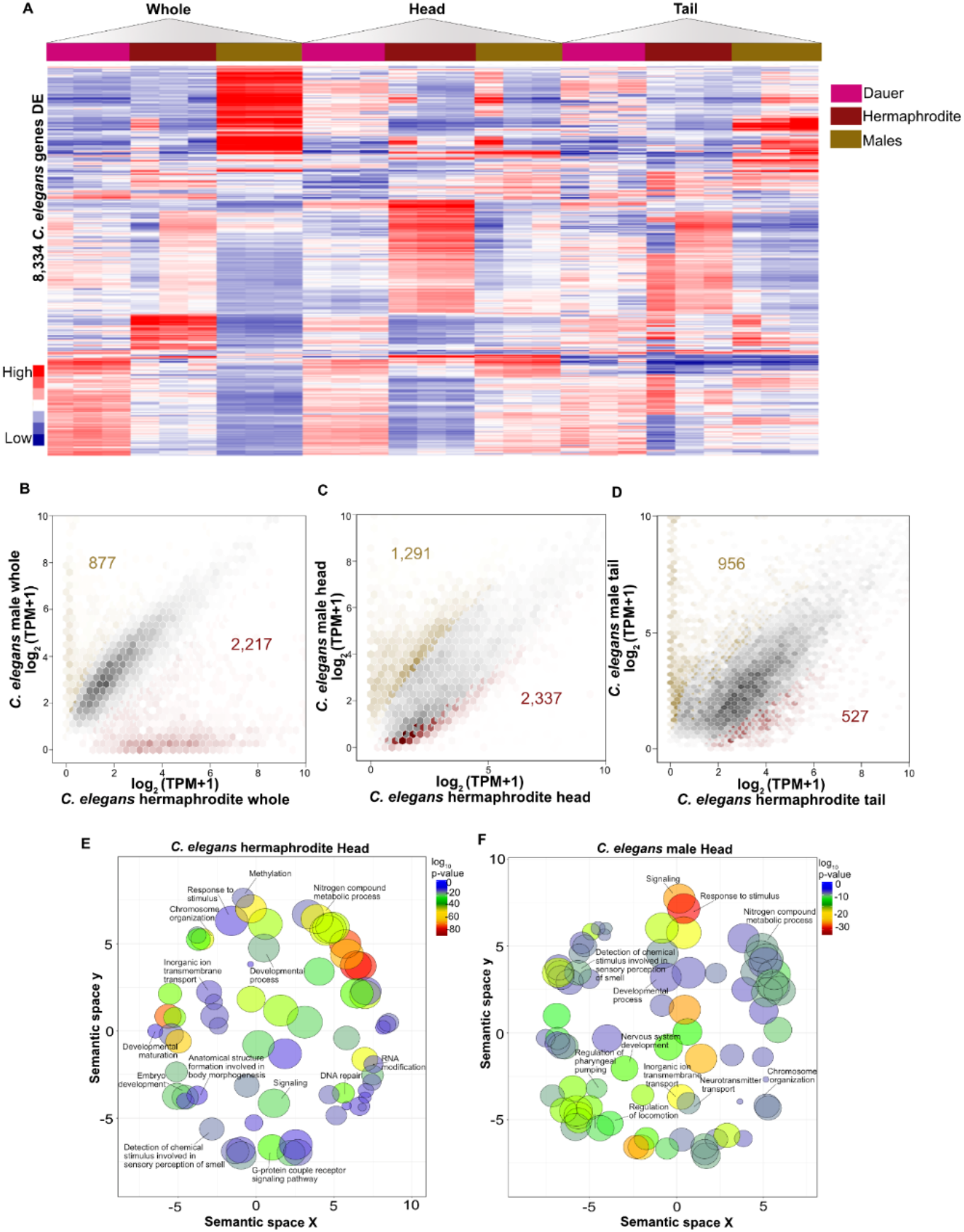
Analysis of differentially expressed genes in *C. elegans* dauers, hermaphrodites, and males. (a) Heatmap showing 8,334 genes with a minimum of 1 transcript per million (TPM). (b, c and d) Hexagonal-bins scatterplots comparing differentially expressed genes between hermaphrodite, hermaphrodite head, hermaphrodite tail, and male, male head, and male tail. Samples originated from hermaphrodites are represented in red and males are represented in dark gold. Each hexagon is divided into six triangles with a range of color intensity to indicate number of observations in each class. (e, f) The list of 3,628 DEGs between hermaphrodite head and male head from panel C were subjected to GO term analysis. The results are shown as REVIGO scatterplots in which GO terms are in a “semantic space” where more similar terms are positioned closer together. The larger the circle indicates a greater number of genes that are included in that GO term.

To characterize region-specific gene activity, we compared young adult heads and tails. First, we compared young adult heads and found that the young male head has 1,291 genes enriched while the young hermaphrodite head has 2,337 genes enriched (Fig. 2C, Table S1). GO terms for young hermaphrodite head were enriched for RNA-DNA processes (*RNA modification, RNA processing, DNA repair, methylation, and chromosome organization*), development (*embryo development, developmental maturation*) and response to the environment (*detection of chemical stimulus involved in sensory perception of smell, response to stimulus, G-protein coupled receptor signaling pathway*) (Fig. 2E). On the other hand, GO terms for males were enriched for neurogenesis (*nervous system development, inorganic ion transmembrane transport, neurotransmitter transport, cell projection organization*) and regulatory components (*regulation of pharyngeal pumping, regulation of stimulus, regulation of locomotion*) (Fig. 2F, Table S2). Hermaphrodite and male heads display neural terms and response to the environment or stimulus as predominant GO terms. These results are as expected since the pharynx is essential for food intake, motor control, isthmus peristalsis, and sensing environmental cues^34^.

Lastly, we examined young adult tails to depict the ontology analysis characteristic of *C. elegans* male tails. We found 527 differentially expressed genes in the hermaphrodite tail in contrast to 956 genes in male tails (Fig. 2D, Table S1). GO terms for hermaphrodite tails were enriched for reproduction (*reproduction, embryo development, gamete generation*), cell cycle (*chromosome organization, the establishment of cell polarity, cell division, chromosome segregation*), and RNA processing (Fig. S2C). GO terms for male tails were enriched for neural responses (*neuropeptide signaling pathway, chemical synaptic transmission, potassium ion transport*), metabolic process (*organic cyclic compound metabolic process, cellular protein metabolic process, phenol-containing compound metabolic process, phosphorus metabolic process*), and detection of chemical stimulus involved in sensory of perception of smell and cilium organization (Fig. S2D, Table S2). For hermaphrodite tails, part of the gonads included in the collection gives rise to GO terms for reproduction. This is in comparison to males which, as expected, have several neural terms which might be due to the cluster of neurons in the tails necessary for reproduction.

Comparative analysis between young adult hermaphrodites and young adult males show significant differences in genes that are enriched in each region. Young whole hermaphrodites are developing all the structures for reproduction, while males are going through intense metabolic processes and neuronal signaling. Young hermaphrodites’ heads and tails show signs of reproductive development while responding to their environment. The same trend also continues with young male heads and tails, in which neurogenesis and metabolic processes make up the vast majority of the GO terms. The enriched GO terms for the neural activity for male head and tail might be due to male-specific neurons such as cephalic male (CEM) neurons and <u>m</u>ystery <u>c</u>ells of the <u>m</u>ale (MCM) neurons in the head^35,36^. These are in addition to mechanosensory and chemosensory neurons in the tail, which are required for detecting the hermaphrodite, locating the vulva and coupling^36–38^. These results show that we detect differences in gene expression between young adult hermaphrodites and young adult males not only when the germline is included but also in separate regions^33^. This data provides a base comparison for equivalent regions in *S. carpocapsae*.

### Transcriptomic comparison of *C. elegans* dauer to young adults

In addition to examining the overall expression of differentially expressed genes (DEGs) in young adults, we also used the RNA-seq data to compare *C. elegans* dauers to young adults. The whole dauer transcriptome has a distinct set of genes that are enriched in comparison to young adult hermaphrodite and young adult males (Fig. 2A). We found 664 genes enriched in dauers when compared to 929 genes in young adult hermaphrodites (Fig. S3A, Table S3). Biological process GO terms for dauers included a response to stimulus, signaling, and cell communication, as well as neural related terms (*signal transduction, neuropeptide signaling pathway, G-protein coupled receptor signaling pathway*) (Fig. S3B, Table S2). Enrichment of genes related to *neuropeptide signaling* included genes such as *flp-1* and *flp-3* in the FMRFamide-like peptides, known to be expressed in the nerve ring and associated with pharyngeal function. They are found to be a conserved gene expressed in dauers between *C. elegans* and other species such as *P.pacificus*^21,29^. GO terms for young adult hermaphrodites were enriched for the molting cycle, immune system, and chemosensory behavior (Fig. S3C). GO terms such as *molting cycle* included genes crucial for to cuticle development and molting including *mlt-4* and *mlt-10*. Other genes included *nhr-23*, whose expression results in arrest and loss of proper molting function, as well as genes responsible for fusion of the cells which form the cuticle, such as *nas-37*, and genes involved in extracellular matrix functions crucial for release of the cuticle, *noah-1* and *naoh-2*^39^. We next compared dauers to young adult males and found 2,761 and 1,001 genes DE respectively (Fig. S3D, Table S3). Biological GO terms for dauers were enriched for dormancy process, regulation of dauer development, dauer entry, and neural related terms (*chemical synaptic transmission, G-protein coupled receptor signaling pathway coupled to cyclic nucleotide second messenger, nervous system development*) (Fig. S3E)(Table S2). Genes essential for dauer entry and dauer development were particularly enriched including *daf-3, daf-4, daf-7, daf-12, daf-38* and *unc-3*^40,41^. Meanwhile, male terms included gamete generation, regulation of growth rate, and phosphorus metabolic process (Fig. S3F). The GO term *gamete generation* included enrichment for genes involved in spermatid development such as *spe-4, spe-8, spe-15, spe-44, try-5*, and *elt-1*, as well as sperm motility and activation including *spe-10* and *swm-1*^42^. Importantly, several GO terms and dauer-specific genes indicate that we indeed collected dauers, while females and males are going through developmental reproductive processes.

Taken together, these results show that young adult hermaphrodites and young adult males share characteristics of developing nematodes^33^. On the other hand, dauers have strong neurogenesis-related terms compared to both young adults. These results support evidence that dauers go through morphological changes in re-wiring of the nervous system, remodeling of dendritic arbors, and glial plasticity^43–45^.

### Transcriptomic comparisons of *S. carpocapsae* young adult males and females

We collected triplicates of the whole, head, and tail of young adult female and young adult males. The *S. carpocapsae* genome has 30,956 genes, of which an average of 26,774 (86.5%) genes were detected in whole young females (at least one mapped read), and 28,830 (93.1%) genes in whole young adult male^46^. For the young female head and tail, we detected 75.6%, and 76.1% of genes were expressed respectively, while for the young male head and tail, we detected 92.1%% and 93.4%. Although there is no validated markers of *S. carpocapsae* region-specific expression, we checked the expression of orthologous genes used above to validate regional gene expression in *C. elegans*. We focused on the one-to-one orthologs and found *flp-1 (L596_g05249)* to have high expression in the head and tail of female, but only in the head of the male (Fig. S4A, B). For females, we found that *pes-8 (L596_g03940)*, expressed in the alimentary system, epithelial system, and gonads, is detected in the whole worm and tail similar to *C. elegans*. Interestingly, *flp-3 (L596_g03561)* is detected the highest in the head and tail similar to *C. elegans*. However, *egl-20 (L596_g05606)* is found in the whole and tail of *S. carpocapsae* as is in *C. elegans* (Fig. S4A). For males, most of the genes did not have one-to-one orthologs. We found that *spe-44 (L596_g014218)*, involved in regulating spermatogenesis, is not expressed at the young adult stage. Lastly, *spe-9 (L596_g018160)*, expressed in the spermatid, is highly expressed in the whole nematode and tail region of *S. carpocapsae*, resembling *C. elegans* expression (Fig. S4B). Although the function of the genes discussed have not been verified in *S. carpocapsae*, characterizing their location provides insights into their function and evolutionary conservation.

Next, we performed a comparative analysis between females and males with a region-specific resolution of gene expression. We found 11,530 genes that were DE in at least one region comparison (Fig. 3A). We established 2,327 genes enriched in young male whole and 1,022 genes in young female whole (Fig. 3B, Table S4). GO terms for young female whole are related to neurogenesis (*neuroepithelial differentiation, mechanoreceptor differentiation*), immunity (*defense response, immune response, immune system process*), cell degradation (*negative regulation of apoptotic process, regulation of autophagy*), several metabolic processes (*lipid metabolic process, indole metabolic process, protein metabolic process*), and regulation of catalytic activity and proteolysis (Fig. S5A, Table S5). GO terms associated with whole young adult male were enriched for sperm maturation (*spermatogenesis, fatty acid transport, acid secretion*), sperm function (*flagellated sperm motility, phosphorus metabolic process, protein dephosphorylation, tyrosine metabolic process*), regulation of biological quality, and protein metabolic process (Fig. S5B). While the young adult males are preparing themselves for reproduction, young females are going through intense metabolic processes and regulation of catalytic activity.

**Figure 3.**
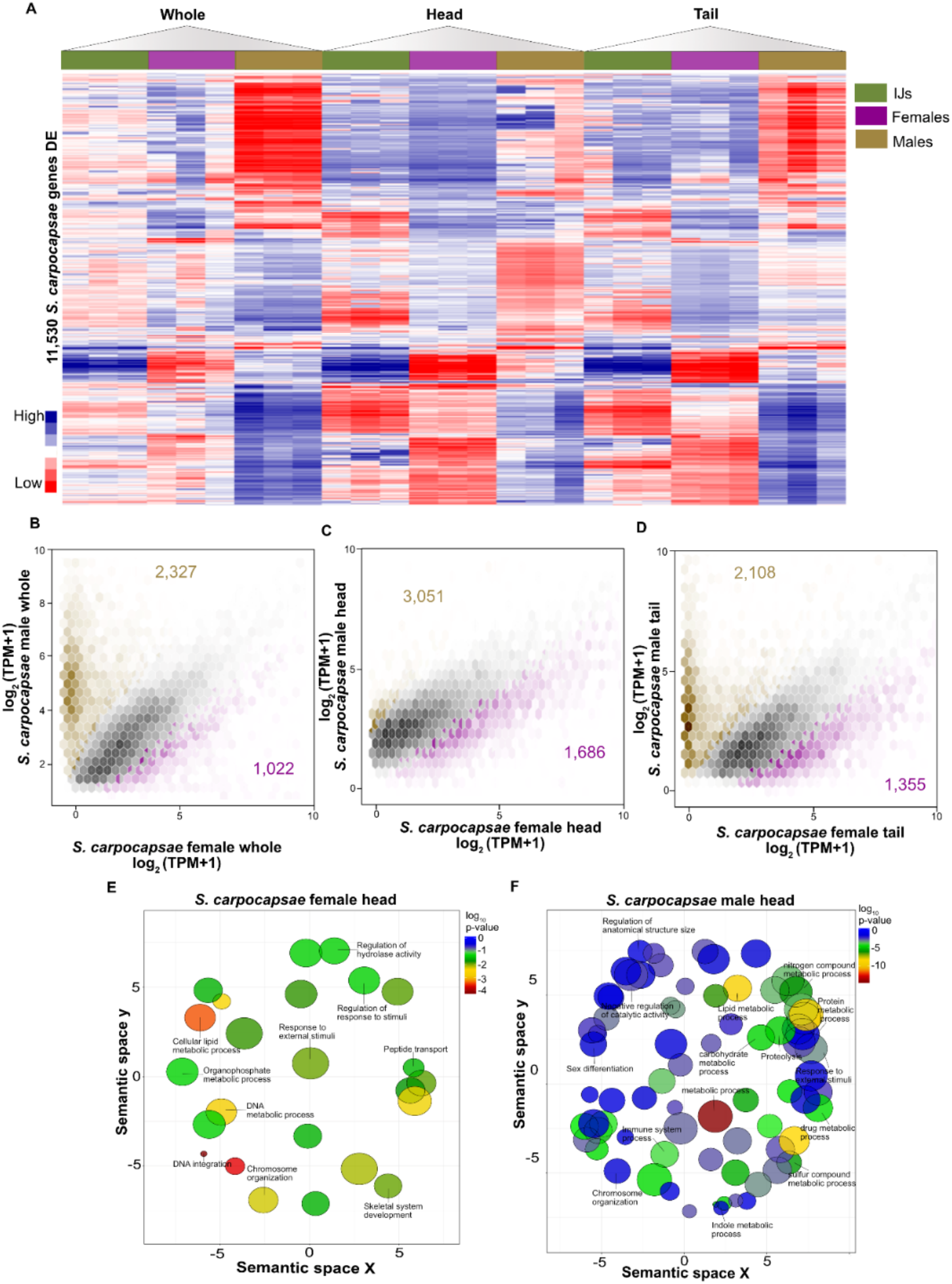
Analysis of differentially expressed genes in *S. carpocapsae* IJs, females, and males. (a) Heatmap showing 11,530 genes with a minimum of 1 transcript per million (TPM). (b, c and d) Hexagonal-bins scatterplots comparing differentially expressed genes between female, female head, female tail, and male, male head, and male tail. Samples originated from females are represented in purple and males are represented in light gold. Each hexagon is divided into six triangles with a range of color intensity to indicate number of observations in each class. (e, f) The list of 4,737 DEGs between female head and male head from panel C were subjected to GO term analysis using REVIGO scatterplots with results displayed in semantic space where more similar terms are positioned closer together.

We then compared heads between young adult females and males. We found 3,051 genes enriched in the young adult female head compared to 1,686 genes in the young male head (Fig. 3C, Table S4). GO terms of biological process for the young adult female head were enriched for regulation of processes such as hydrolase activity and response to stimuli. They also included DNA processes (DNA metabolic process, DNA integration, chromosome organization), as well as cellular lipid process, response to external stimuli, and skeletal system development (Fig. 3E). On the other hand, biological process GO terms for the young male head are related to different types of metabolism, including sulfur compound, drug, carbohydrate, organophosphate, lipid, nitrogen compound, and indole, additionally to negative regulation of catalytic activity, sex differentiation, and regulation of anatomical structure size (Fig. 3F, Table S5). Female and male *S. carpocapsae* are both developing structures, indicating that development is not complete yet. While females are responding to their environment, males are undergoing intense metabolic processes.

We next identified the genes that were enriched in the tail of females and males. Females have 1,355 genes enriched compared to 2,108 genes in males (Fig. 3D, Table S4). Biological GO analysis for the female tail suggested terms related to positive regulation of Notch signaling, lipid storage, sensory perception of sound, and neural related terms (neuropeptide signaling pathway, neuroepithelial cell differentiation) (Fig. S5C). The vast majority of biological process GO terms for the young male head were related to reproduction including *spermatogenesis, flagellated sperm motility, tyrosine metabolic process, phosphorus metabolic process*, and *acid secretion*. They also included ribosome biogenesis, response to a stimulus, and response to growth factors (Fig. S5D, Table S5). The female tails are sensing their environment while developing. In contrast, males are largely preparing themselves at the young adult stage for mating.

Our comparison of *S. carpocapsae* young adult females to males revealed that females are developing while males are forming reproductive structures. The whole females and tails had several terms related to neuron formation and differentiation; interestingly, tail projections found in the first generation of females in the *Steinernematidae* family have yet to be characterized and might play a role in neuronal activity detected in GO terms^47^. As expected, the female head is regulating and responding to external stimuli, which agrees with the function of the pharynx^34^. In the whole female, we also detect several metabolic processes and immunity related terms which may be indicative of the intestine tissue^48^. In contrast, young adult males are developing the structures and sperm needed for mating. The whole nematode, head, and tail have GO terms related to reproduction. Lastly, the male head has several metabolic processes related to hypodermis tissue^48^.

### Transcriptomic comparison of *S. carpocapsae* IJs to young adults

We then compared IJ whole nematode to young adult male and female to acquire insights into reproductive structures, morphology, and behavior. In whole IJs we detected 28,753 (92.9%) genes of which a subset of genes shared similar expression to females and another subset to males (Fig. 3A). We then performed a comparative analysis of whole IJs to whole young adult females and found 2,566 and 2,015 DEGs respectively (Fig. S6A, Table S6). Biological process GO terms for IJs had several related to neural activity (*synaptic vesicle cycle, neurotransmitter transport, chemical synaptic transmission, neuropeptide signaling pathway*), regulatory processes (*regulation of mesoderm, regulation of metabolic process, regulation of cellular process*), signaling, and response to odorant (Fig. S6B, Table S5). Of the 2,566 differentially expressed genes in IJs compared to females there were approximately 885 of those that were one-to-one orthologs with *C. elegans*. Genes involved in nervous development including daf-1(L596_g019493), and genes involved in the Wnt signaling pathway (*lin-17(L596_g09403), lin-44(L596_g010782), pry-1(L596_g026285), cfz-2(L596_g012382)*). Additionally, some of the orthologs included genes known to be involved in *C. elegans* dauer entry such as *daf-10 (L596_g05248), daf-7 (L596_g018475*), and *daf-16 (L596_g010238*)^30,40,49,50^. GO terms for young adult females were enriched for metabolic process (*phosphorus, protein, nitrogen*), as well as embryo development, reproduction, developmental process and negative regulation of catalytic activity (Fig. S6C) (Table S5). Within the 2,015 differentially expressed genes in young adult female, 656 one-to-one orthologs included genes annotated in *C. elegans* to be involved in nematode larval development and embryogenesis in such as *mlc-4* (L596_g025459), *arx-2* (L596_g027714) as well as genes in *C. elegans* known to be involved in ovulation and gametogenesis such as *deb-1* (L596_g04265) and *unc-15* (L596_ g06843)^51,52^.

Next, we compared IJs to young adult males. We found 3,385 genes enriched in IJs compared to 3,662 genes in young adult males (Fig. S6D, Table S6). GO terms for IJs included several metabolic processes (*nitrogen compound, cellular aromatic compound, drug*), sleep, response to a stimulus, negative regulation of behavior, cell communication, and neural terms (*nervous system development, chemical synaptic transmission, neuropeptide signaling pathway*) (Fig. S6E, Table S5). Of the 3,385 genes found to be differentially expressed in IJs, approximately 1,221 of those genes have one-to-one orthologs with *C. elegans*. Within those 1,221 orthologs consistent with the female comparison were genes annotated in *C. elegans* to be involved in dauer entry and development such as *daf-37* (L596_g01775), *daf-38* (L596_g04980), *daf-9* (L596_g018548), *daf-7 (L596_g18475)* and *daf-16 (L596_g010238)*. Additionally present were orthologs related to neuropeptide signaling pathways, including a number of genes related to two major class of neuropeptides known in *C. elegans* as FMRFamide-like neuropeptides (*flp-1 (L596_g05249) flp-3* (L596_g03561), *flp-7*(L596_g03276)) some of which are known to be highly conserved in nematodes, and neuropeptide-like protein (*nlp*) genes (*nlp-19*(L596_g029787), *nlp-3*(l596_g03984), *nlp-13*(L596_g027854) and *nlp-21*(L596_g018927))^29,53,54^. Meanwhile, young adult male terms included developmental maturation, germ cell development, chemical synaptic transmission, and G-protein coupled receptor signaling pathway, in addition to several metabolic processes such as tyrosine, organonitrogen, phosphorus, and nitrogen (Fig. S6F). Of the 3,662 genes found to be differentially expressed in young adult males there were only about 537 conserved one-to-one orthologs with *C. elegans* genes annotated for being involved spermatid development including *spe-6* (L596_g016164), *spe-10* (L596_g017136) *spe-4* (L596_g015636) and for involvement in embryo and male tail development *lin-41* (L596_g07872), *spg-7* (L596_g09294), and *ran-3* (L596_g07335)^42,55^. The terms for IJs were heavily geared towards neural related processes, while females and males are going through development, and several metabolic processes are involved for both young adults.

In summary, these results show that IJs are enriched for neural activity which included several conserved neuropeptides and dauer entry and development genes when compared to young adult females and males. Additionally, despite having fewer differentially expressed genes compared to young adult males and more compared to females, overall IJs had a consistently higher percentage of genes that were one-to-one orthologs with *C. elegans* within the genes that were differentially expressed in each comparison. The enriched activity during this stage correlates with host-seeking behavior and sensory responses^56^. On the other hand, females and males are growing and developing reproductive structures. Each has several metabolic processes, some of which like phosphorus metabolism are important for reproduction, while other compound metabolisms such as nitrogen are related to the hypodermal tissue.

### *S. carpocapsae* young adult female reveals contrasts in gene expression of metabolic processes, neurogenesis and development compared to *C. elegans* young adult hermaphrodite

We compared whole, head, and tail of *C. elegans* young adult hermaphrodite and male to the equivalent of *S. carpocapsae* young female and male. The transcriptomic comparisons provided insight into the similarities and differences of signaling pathways and developmental processes shared between these two distantly related species. We compared across species between corresponding sex (hermaphrodite versus female, male versus male) and region (head and tail). First, we identified 6,136 one-to-one orthologs between *S. carpocapsae* and *C. elegans (Table S7*). Then, we took orthologous genes and performed a comparative analysis between equivalent sex and region and found 2,732 genes were enriched in at least one comparison across species (Fig. S7A).

Interestingly, 115 genes were lowly expressed (less than 1 TPM) or not expressed (0 TPM) in any of the regions (Table S8). GO terms for the 115 genes were enriched for DNA and RNA activity, developmental processes, and neurogenesis (Fig. S7B). Because the 115 genes were lowly or not expressed at all, we hypothesized that we would detect the expression of these genes across previously sequenced 20 developmental stages would be higher than in adults^46^. Indeed, the expression of the 115 genes is detected at some point between zygote to 4-cell, 8-cell to 3-fold, and L1 to 15 hours^57^ (Fig. S7C). We conclude that this gene set, which is shared between both species, is highly conserved and likely crucial for developmental processes from zygote to first larval stage. This brings an important point that genes differentially expressed in a region-specific manner in both species of *S. carpocapsae* and *C. elegans* will provide insight into the developmental processes that are conserved in these two distantly related species.

First, we investigated each whole nematode and performed region-specific comparative analysis between *S. carpocapsae* young female and male and *C. elegans* young hermaphrodite and male to understand similarities and differences across each region. Regio-specific analysis provides high-resolution of gene expression without amplifying genes important for gonad development. Differential expression analysis uncovered 766 genes enriched in *S. carpocapsae* whole young female nematodes and 1,097 genes in whole *C. elegans* young hermaphrodite nematodes (Fig. 4A, Table S9). GO analysis revealed terms that are similar between *S. carpocapsae* and *C. elegans* such as locomotion, regulation of response to a stimulus, and developmental processes (Fig. S8A, Table S10). The shared GO terms stem from different sets of genes being expressed at the equivalent stage and region but are used for a similar function. The shared GO term “developmental processes” in *S. carpocapsae* young females represents 113 genes which encompass pathways such as Wnt signaling (*siah-1, kin-20*)^58,59^, and several genes categorized in angiogenesis (*jnk-1, sem-5, mpk-1, let-60, pxl-1*)^60–64^. The genes that fell in the category of angiogenesis are important for *C. elegans* vulva development^65^. The Wnt signaling genes are known to be part of linker cell type death^66^ (*siah-1*) and seam cell development^58^ (*kin-20*). On the other hand, we found 196 genes in *C. elegans* young hermaphrodite for the “developmental processes” term. The pathways included Wnt signaling^50^ (*ssl-1, egl-30, mom-2, lin-17, lin-44, cdh-3, cfz-2, lit-1, fmi-1*) in addition to genes categorized in angiogenesis^67,68^ (*vps-34, aap-1*). The vast majority of the Wnt signaling genes are part of vulva development, while the genes in angiogenesis are important for growth and lifespan determination^65^. Because gonad development takes most of the tissue in a whole nematode, we expected many genes to be related to vulva development.

**Figure 4.**
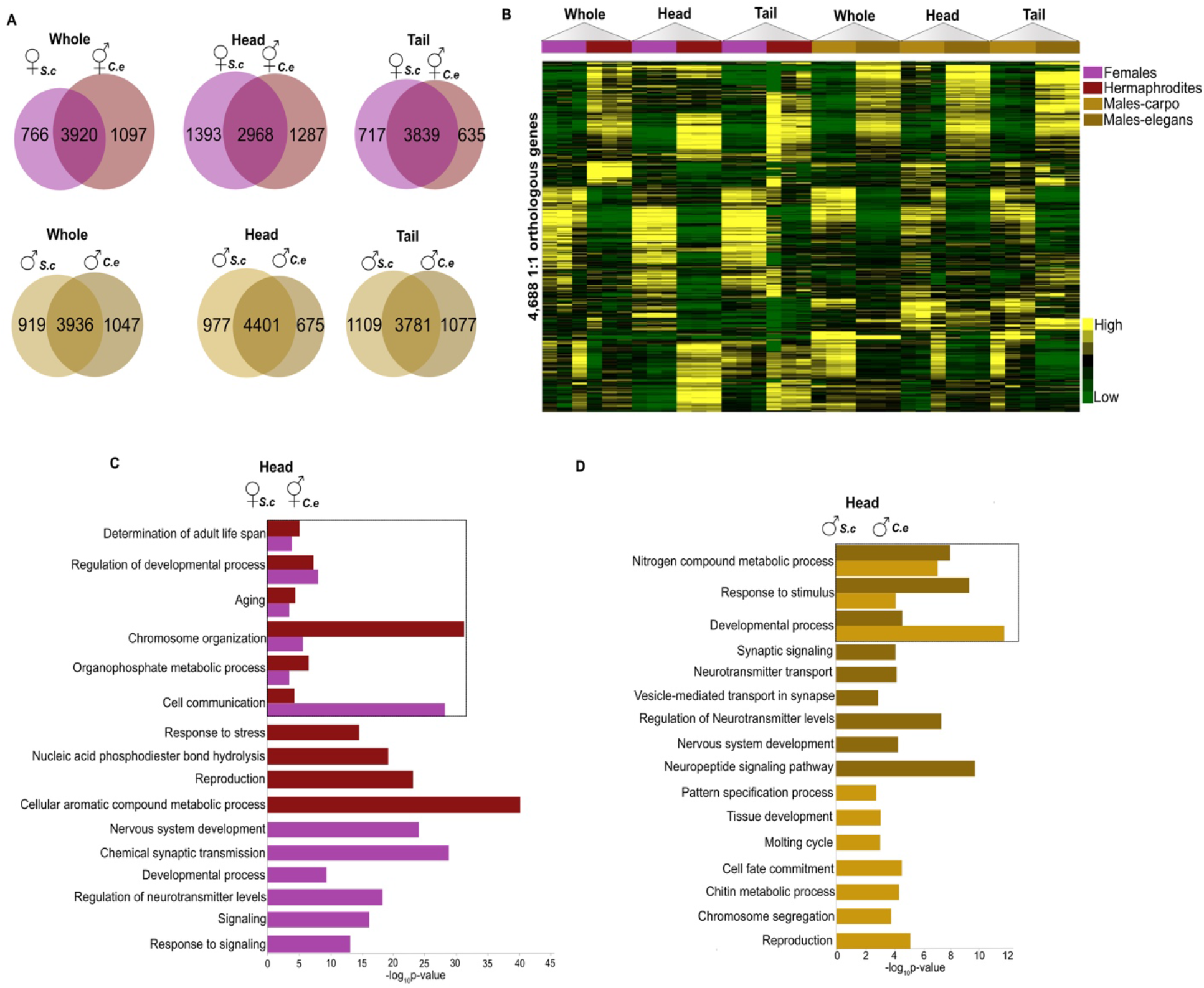
Differential gene expression analysis of one-to-one orthologs between *Steinernema carpocapsae* males and females and *Caenorhabditis elegans* males and hermaphrodites. (a) Venn diagrams showing differentially expressed genes between *S. carpocapsae* and *C. elegans* whole worms, heads and tails. Tissues were compared between *S. carpocapsae* female (purple) and *C. elegans* hermaphrodite (red) and *S. carpocapsae* male (light gold) and *C. elegans* male (dark gold). (b) Heatmap showing 4,688 differentially expressed one-to-one orthologs in at least 1 tissue comparison between *S. carpocapsae* and *C. elegans*. (c, d) Gene ontology analysis of DEGs between *S. carpocapsae* female and *C. elegans* hermaphrodite heads, and *S. carpocapsae* male and *C. elegans* male heads from panel A. Colors reflect the nematode stage and the bars are indicative of the P-value for the associated GO terms. Black box indicates shared GO terms.

Although the analysis between species had a few shared terms, *S. carpocapsae* whole young female nematode and *C. elegans* whole young hermaphrodite nematode had their own sets of unique GO terms. *S. carpocapsae* whole young female GO terms were related to metabolic processes such as phosphorus, lipoprotein, lipid, and organophosphate (Fig. S8A). In addition to these, we found functions relating to tissue development (*mesoderm development, animal organ development*), and sensory perception of a chemical stimulus. On the other hand, the 1,097 genes enriched in *C. elegans* young hermaphrodites whole revealed functions relating to neuronal signaling (*neuropeptide signaling pathway, synapse organization, synaptic signaling*) and growth (*tissue development, regulation of growth, molting cycle* and *cuticle development*) (Fig. S8A). *S. carpocapsae* terms are enriched for metabolic processes, in contrast with *C. elegans*, which have intense neuronal activity.

We then compared the heads of females and hermaphrodites to obtain a more explicit depiction of region-specific gene expression. Head tissue of *S. carpocapsae* young females had 1,393 enriched genes and *C. elegans* young hermaphrodites had 1,287 enriched genes (Fig. 4A, Table S9). After the DE analysis, we looked into shared GO terms and found several, such as organophosphate metabolic process, cell communication, chromosome organization, regulation of developmental process, aging, and determination of adult lifespan (Fig. 4C). We further looked into “aging” because of its extensive study in pathways such as insulin/insulin-like growth factor-1 signaling (IIS), which first established lifespan-regulating signaling pathway in *C. elegans*^69^. The “aging” GO term represented 35 genes in *S. carpocapsae* female heads, which encompassed pathways such as TGF-beta dauer pathway (*daf-14, daf-7*), famous for longevity in *C. elegans*, and the gene *ral-1* which is critical for cell-cell communication via exosomes^41,70^. We also found 36 genes associated with “aging” in *C. elegans* hermaphrodite heads, which included the Insulin/IGF pathway kinase B signaling cascade (*daf-18, aap-1*) and *vps-34*, all which are essential for nematode longevity^71,72^. Apart from shared GO terms, *S. carpocapsae* young female heads had unique terms associated with the nervous system (*nervous system development, chemical synaptic transmission, neurotransmitter levels*), metabolic activity (*phosphorus metabolic process, organonitrogen compound metabolic process*) and terms relating to response to stimulus and reproductive behavior. GO terms enriched in the *C. elegans* young hermaphrodite head showcased functions relating to *the determination of adult lifespan*, reproduction, DNA metabolism, response to stress, detection of chemical stimulus involved in sensory perception of smell and detection of chemical stimulus. These results show that at equivalent stages, both species share different genes related to aging, and the *S. carpocapsae* female head has a more pronounce metabolic and neurogenesis activity in contrast to the *C. elegans* hermaphrodite head, which is sensing the environment and determining its lifespan.

We next contrasted the enriched genes in the tail region of *S. carpocapsae* female and *C. elegans* hermaphrodite to understand the differences in gene expression that correlates with tail function. Tail function in hermaphrodite is unspecialized, but all the neurons in this region have been identified^73^*. S. carpocapsae* young female tails expressed 717 genes compared to 635 genes in *C. elegans* young hermaphrodite tails (Fig. 4A, Table S9). GO biological process terms found to be shared between tail stage-enriched genes included terms related to metabolic process (*organophosphate, cellular aromatic compound, phosphorus*), vesicle-mediated transport, regulation of biological quality, and sensory perception of chemical stimulus (Fig. S8B). Among the shared terms in *S. carpocapsae* female tail, we found “regulation of signaling” to have 32 genes, which encompassed pathways such as fibroblast growth factor (FGF) and Wnt signaling pathway. The FGF pathway had genes such as *sos-1*, important for RAS mediated developmental signal, *tpa-1*^74^, involved in axon regeneration, and *ptp-2*^75^, necessary for muscle and vulva development. The Wnt pathway had two genes: *egl-30*, involved in vulva development, and *pry-1*, important in lipid metabolism^76^. However, in *C. elegans*, young hermaphrodite tails “regulation of signaling” represented 39 genes, including *sem-5*, an epidermal growth factor part of the FGF signaling pathway, and *kin-20*, a circadian clock gene part of the Wnt signaling pathway^61,77^. We lastly looked at the terms unique to *S. carpocapsae* female tail that displayed functions relating to metabolic processes (*phosphate, lipid, phosphorus*), and cell signaling (*Ras protein signal transduction, intracellular transport, regulation of cell communication*). In contrast, different terms for *C. elegans* hermaphrodite tail were associated with locomotion, regulation of locomotion, regulation of signaling, reproduction, and neurogenesis (*nervous system development* and *synapse organization*). These results suggest that *S. carpocapsae* female tail is enriched for metabolic processes while *C. elegans* hermaphrodite tail had expected activity such as locomotion and neurogenesis.

The comparison of *S. carpocapsae* female to *C. elegans* hermaphrodite reveals contrasts in metabolic processes, neurogenesis, and development. *S. carpocapsae* female whole, head, and tail are enriched for metabolic processes, including lipoprotein and lipid protein, which in *C. elegans* has been extensively studied concerning the regulation of fat and energy metabolism as well as starvation^78,79^. Both species shared the GO term “developmental processes,” which encompassed several genes essential for *C. elegans* vulva development^53^. Another shared GO term was related to “aging,” with numerous genes that have been identified in *C. elegans*, which regulates its lifespan^80^. The genes that are differentially expressed in both species and share a common function present interesting future questions about core genes involved in development and aging in distantly related species.

### *S. carpocapsae* young adult male reveals contrasts in gene expression of morphogenesis, neural signaling and development compared to *C. elegans* young adult males

Comparative transcriptional analysis was also carried out between *S. carpocapsae* young males and *C. elegans* young males with an emphasis on the developmental process to characterize gene expression patterns. The young adult whole worm tissue comparison revealed 919 genes enriched in *S. carpocapsae* and 1047 genes enriched in *C. elegans* (Fig. 4A, Table S9). The GO analysis had few shared terms between species such as developmental process, regulation of locomotion, regulation of response to a stimulus, and metabolic processes such as *nitrogen compound and organic substance* (Fig. S9A, Table S10). We further looked into the genes differentially expressed in the term “developmental process” to acquire insights into biological differences at the early adult male stage. For *S. carpocapsae*, the term represented 148 genes involved in several pathways. We found genes such as *dcap-1*, predicted to be a decapping enzyme important for adult life span determination and nematode larval development^81^. We also looked into pathways such as PDGF-signaling (*gsk-3, let-60, rsks-1*), necessary for neural and glial development^82–84^.

In contrast, for *C. elegans*, the term “developmental process” had 151 genes. Interestingly, the PDGF-signaling pathway was also present in the ontology analysis, but with a different set of genes (*ras-1, ets-4, vps-34*), from which we can reiterate the idea of similar pathways being conserved but used differently by each nematode stage. We then focused on unique GO terms for each male species. For *S. carpocapsae* whole young adult male, GO terms were associated with reproduction (*reproduction, Phosphorus metabolic process, regulation of reproductive process*), morphogenesis (*mesoderm development, animal organ development, tissue development, chitin metabolism*), and metabolic activity (*regulation of hydrolase activity, organophosphate metabolic activity* (Fig. S9A). On the other hand, GO analysis of the genes enriched in *C. elegans* whole young adult males indicated functions relating to the nervous system (*nervous system development, synapse organization*), neuronal signaling (*neuropeptide signaling pathway, inorganic ion transmembrane transport, regulation of neurotransmitter levels, chemical synaptic transmission*), and *reproductive behavior* (Fig S9A). The comparison of both whole young adult males revealed that they are developing structures for the nervous system, locomotion, and reproduction. Despite this similarity, *S. carpocapsae* also has several metabolic activities, conversely to *C. elegans*.

We proceeded to a comparative analysis on the young adult male heads between *S. carpocapsae* and *C. elegans* to further characterize a head-specific transcriptional profile. Expression comparisons revealed 977 genes enriched in the *S. carpocapsae* young adult male head, while 675 genes were enriched in the *C. elegans* young adult male head (Fig. 4A) (Table S9). After the DE analysis, we looked at shared biological process GO terms and found several such as *developmental process*, *response to stimulus* and metabolic processes for *nitrogen compound, organic cyclic compound, macromolecule*, and *organic substance*. Again, we looked at the shared GO term “developmental process” for *S. carpocapsae* and *C. elegans*. For *S. carpocapsae* young male heads the term encompassed 155 genes involved in several pathways, such as genes part of Notch signaling, *sel-12*, important for mesoderm patterning and muscle function, and *lin-22*, involved in the patterning of the peripheral nervous system in *C. elegans*^85,86^. Another well-studied pathway that came up was the Wnt signaling pathway (*egl-20, lin-44, cdh-3, siah-1, cwn-1, kin-20, spe-6*) which, in *C. elegans*, is an crucial morphogen that provides developing tissue positional information and specify cell polarity and migration^30,87^. For *C. elegans*, the term “developmental process” had 92 genes involved in pathways like integrin signaling (*unc-97, emb-9, vps-34, lam-2*), important for cell-matrix interactions and a focus of studies of cell invasion^88,89^. Apart from the several shared GO terms, *S. carpocapsae* young adult male heads had different terms associated with development and morphogenesis (*tissue development, molting cycle, cell fate commitment, chitin metabolic process, pattern specification process*), reproduction and chromosome segregation (Fig. 4D). *C. elegans* young male heads GO terms suggested functions relating to nervous system development, neuronal synaptic signaling (*neurotransmitter transport, vesicle-mediated transport in the synapse, regulation of neurotransmitter levels, neuropeptide signaling pathway and synaptic signaling*), and metabolic processes (*organophosphate, organonitrogen*) (Fig. 4D, Table S10). The male heads are finishing development indicated by several genes in the Wnt pathway. Additionally, *S. carpocapsae* is also preparing for reproduction, while *C. elegans* is focused on the nervous system development and neural signaling.

Lastly, an identical comparative analysis was performed between the tail regions to acquire insight into male specific structure and morphology. We identified 1,109 genes enriched in *S. carpocapsae* young adult male tails and 1,077 in *C. elegans* young adult male tails (Fig 4A). GO terms shared between the young adult male tails included *developmental process, detection of chemical stimulus involved in sensory perception of smell*, and *cell component organization* (Fig S9B). Identically to the previous comparisons, we looked into the shared GO term “developmental process.” In *S. carpocapsae*, we found it to represent 181 genes and include several pathways such as Wnt signaling with *cfz-2, part of the frizzled family*, and *cwn-2*, part of the five Wnts, and both are involved in establishing cell polarity, cell migration, asymmetric cell division, and determining cell fate^90,91^.

Additionally, we found *swsn-1*, part of the core SWI/SNF complex and vital for determining *C. elegans* lifespan, and *spe-6*, which has serine/threonine kinase activity involved in spermatid differentiation^92,93^. For *C. elegans* the “developmental processes” GO term encompassed 186 genes and numerous pathways including muscarinic acetylcholine receptor (*egl-30, unc-17, tpa-1*), which is an essential pathway for responding to pathogens, neuromuscular function, and is involved in antimicrobial peptide production expressed in the tail^94–96^. After the analysis of the shared GO term “developmental process,” we then looked at GO analysis of the genes enriched in *S. carpocapsae* young male tail. We found terms relating to morphogenesis (*anterior/posterior pattern specification, embryo development and developmental processes*), DNA packaging (*chromosome organization, chromatin organization, chromosome segregation*), and reproductive processes (*phosphorus metabolic process, reproduction*) (Fig. S9B). Meanwhile, genes enriched for the *C. elegans* young male tails were found to be involved in functions relating to locomotion, nervous system development, cell organization and neuronal communication such as chemical synaptic transmission, G-protein coupled receptor signaling pathway, coupled to cyclic nucleotide second messenger, vesicle-mediated transport in synapse, and neurotransmitter transport (Fig. S9B). As shown above, both species are differentiating their tail structures and using the Wnt signaling pathway. *S. carpocapsae* male tails show signs of morphogenesis and reproduction, while *C. elegans*’ has intense neuronal activity.

Our gene expression comparison between young adult males of distantly related species reveals potential differences in morphogenesis, reproduction, and neuronal activity. We investigated the region-specific GO term “developmental process” and detected the presence of the Wnt pathway in *S. carpocapsae*, which correlates with the importance of Wnt for development in *C. elegans*^97^. We also detected gene expression differences related to reproductive processes present in *S. carpocapsae* but not in *C. elegans*. Conversely, *C. elegans* has several neural and developmental processes and neural communication-related terms when compared to *S. carpocapsae*. The enrichment of go terms in *C. elegans* for neural activity raises the question of the evolutionary conservation of neuronal genes among distantly related nematodes.

### *C. elegans* dauer transcriptome compared to *S. carpocapsae* IJ

We proceeded to a transcriptomic comparison of *C. elegans* dauer and *S. carpocapsae* IJ with region-specific resolution. First, we investigated the expression of the 6,136 one-to-one orthologs by implementing K-means clustering, in which cluster 1 and cluster 2 are genes grouped by species, while cluster 3 displays the shared set of genes across the two species (Fig. 5A). We then explored the differences between whole *S. carpocapsae* IJs and *C. elegans* dauers to capture changes in gene transcription that also occurs throughout the body. *C. elegans* whole dauers displayed 863 enriched genes, while *S. carpocapsae* whole IJs revealed 1,289 enriched genes (Fig. 5B) (Table S11). IJs and dauers shared GO terms such as tissue development, phosphorus metabolic process, and developmental process. Within tissue development, *S. carpocapsae* IJs were enriched for 25 genes, while *C. elegans* dauers showed expression of 23 genes (Fig. S10A) (Table S12). Although IJs and dauers shared GO terms, we found different sets of genes involved in similar functions. For example, integrin signaling, a pathway found under “tissue development” for both species, has different sets of genes. *S. carpocapsae* IJs utilize *arx-7*, expressed in the hypodermis and linker cell, *rap-2*, a GTPase necessary for collagen and cuticle development, and *src-1*^98^, a signaling tyrosine kinase required for cell migration. In contrast, dauers express *epi-1*, which is necessary for epithelialization of various tissues, *arx-4*, part of the Arp2/3 complex necessary for associative learning, and *lam-2*, which is a laminin important for regulation of locomotion^99–101^. We further investigated different GO terms for *S. carpocapsae* whole IJ and found terms related to pseudouridine synthesis, RNA modification, RNA metabolic processes, mesodermal cell fate specification, tissue morphogenesis, nucleoside monophosphate biosynthetic processes, and organic substance biosynthetic process (Figure. S10A). In contrast, biological processes GO analysis of *C. elegans* dauers showed genes related to motor neuron axon guidance, chemotaxis, *axonogenesis*, axonal fasciculation, neuron development, neurogenesis, *developmental process*, and dauer entry (*entry into diapause, dauer larval development*). Overall, the *S. carpocapsae* IJ is going through RNA activity and tissue specification, contrary to the *C. elegans* dauer, which is enriched for neuronal activity.

**Figure 5.**
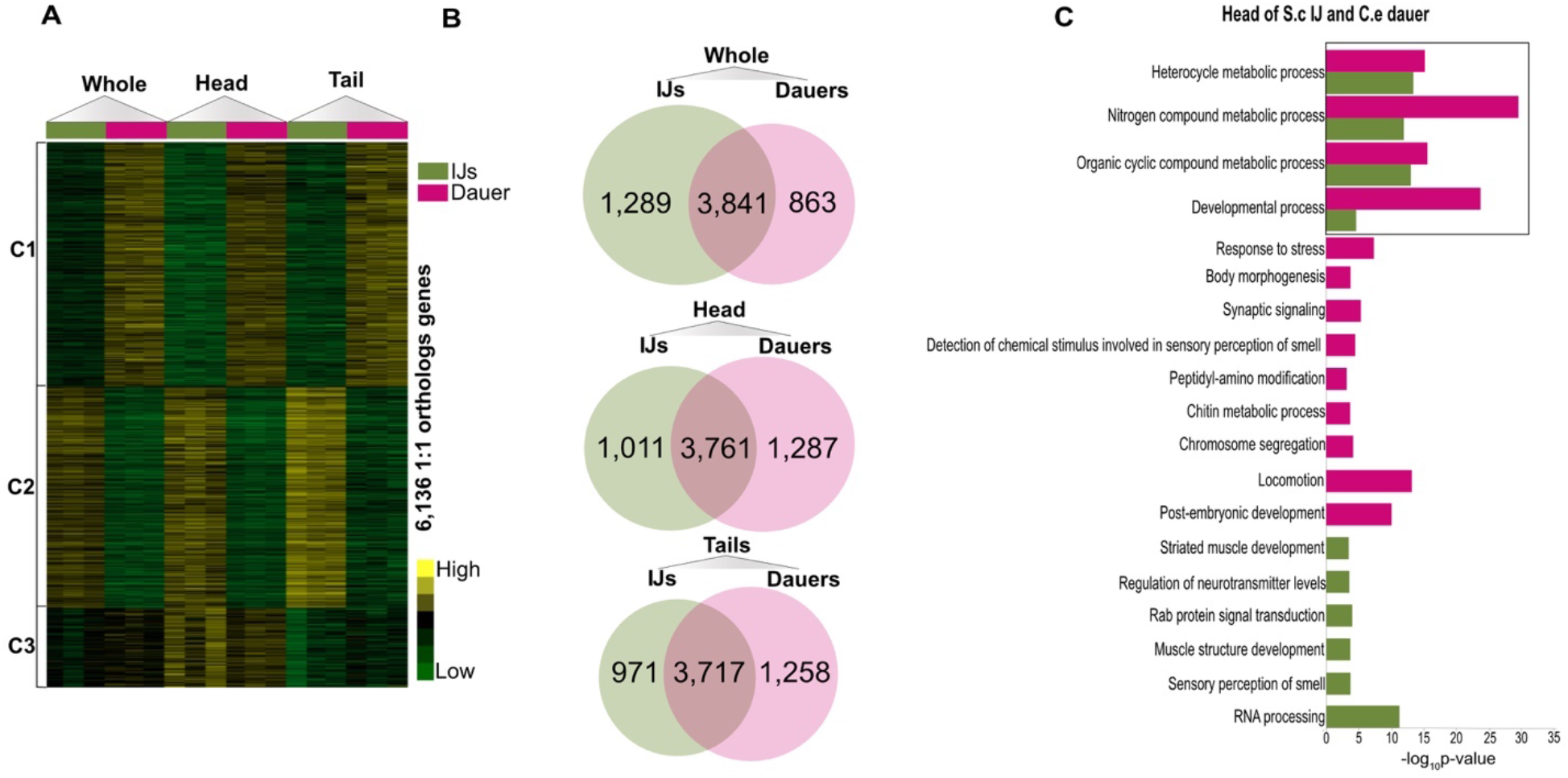
Differential gene expression analysis of one-to-one orthologs between whole, head and tail of *Steinernema carpocapsae* IJ to *Caenorhabditis elegans* dauer. (a) Heatmap showing K-means of 6,136 one-to-one orthologs. (b) Venn diagrams showing differentially expressed genes between *S. carpocapsae* IJ and *C. elegans* dauer whole worms, heads and tails. Regions were compared between *S. carpocapsae* IJ (green) and *C. elegans* dauer (pink). (c) Gene ontology analysis of DEGs between *S. carpocapsae* IJ head (green)and *C. elegans* dauer head (pink). The bars are indicative of the P-value for the associated GO terms. Black box indicates shared GO terms.

The heads of both species were then evaluated to determine region-specific gene expression. We compared *S. carpocapsae* IJ head to that of *C. elegans* dauers, and we saw 1,011 genes enriched in IJs and 1,287 enriched in dauers (Fig. 5B) (Table S11). GO terms shared between both species were heterocycle metabolic process, nitrogen compound metabolic process, organic cyclic compound metabolic process, and developmental process. In *S. carpocapsae*, further investigation of the shared “developmental process” GO term, which represented 128 genes, revealed pathways relating to cytoskeletal regulation by Rho GTPase (*arx-7, arx-6, unc-60, pfn-3*) and dopamine receptor mediated signaling (*gsp-2, snb-1*). In *C. elegans*, the “developmental process” encompassed 233 genes, which in the p53 pathway (*vps-34, atl-1, sir-2.1*) are essential for cell division, phagosome maturation, and dauer larval development^67,102,103^. We further looked into different GO terms for *S. carpocapsae* IJ heads, which included synaptic vesicle exocytosis, neurotransmitter secretion, neurotransmitter transport, signal release, striated muscle cell development, and muscle structure development (Fig. 5C) (Table S12). Conversely, *C. elegans* dauer heads yielded terms relating to the regulation of sensory neuron axon guidance, regulation of cell differentiation, neurogenesis, detection of chemical stimulus involved in sensory perception of smell, and positive regulation of multicellular organism growth. Notably, high-resolution gene expression of both *S. carpocapsae* IJ and *C. elegans* dauer heads displayed terms enriched for neuronal activity and development.

Following the analysis of IJ and dauer heads, we next compared gene expression in their tails. Examining the differentially expressed genes, we found 971 genes enriched in *S. carpocapsae* IJ tails and 1,258 genes in *C. elegans* dauer tails (Fig. 5B) (Table S11). GO terms that were enriched in both *S. carpocapsae* IJ tails and *C. elegans* dauer tails include phosphorus metabolic process, muscle structure development, and organophosphate metabolic process (Fig. S10B) (Table S12). The GO term “muscle structure development” involved 17 genes in *S. carpocapsae* and 15 genes in *C. elegans*. This GO term includes the integrin signaling pathway, with *unc-97, emb-9, pat-3*, and *pat-4* being expressed in *S. carpocapsae* and *let-60* expressed in *C. elegans*. We then looked at GO terms unique for *S. carpocapsae* IJ tails, which included anatomical structure development, developmental process, detection of stimulus, and hemidesmosome assembly. *C. elegans* dauer tails yielded GO terms involved in dauer larval development, positive regulation of developmental growth, motor neuron axon guidance, *axonogenesis, chemotaxis, response to an external stimulus*, and sensory perception. Overall, in *S. carpocapsae* IJ and *C. elegans* dauer tails, several GO terms were describing metabolic and developmental processes.

In summary, the comparison between *S. carpocapsae* IJ to *C. elegans* dauer reveals differences for genes involved in growth and neurogenesis. *S. carpocapsae* IJs whole and tail had several terms related to tissue growth and development, while in *C. elegans*, the same tissues were enriched for neuronal signaling and activity. As expected, both species heads shared terms for neuronal activity and growth. Although *C. elegans* has been an intensively characterized model for neuronal function, much less is known about distantly related nematodes, with the primary assumption that this would be highly conserved among clades 9-12^104,105^. However, *S. carpocapsae* IJs have 40% more ventral nerve cord (VNC) neurons than *C. elegans*^106^. This disparity in the numbers of VNC neurons in addition to the absence of neural terms found between one-to-one orthologs can be a point of interest to identify genes that are neural specific for *S. carpocapsae* IJ.

## Discussion

Here we exploited the morphological similarities between *C. elegans* and *S. carpocapsae* to understand region-specific gene expression. From both species, we collected and sequenced heads and tails as well as the whole worm. We chose two stages of development in *S. carpocapsae*, young adults and IJs, to compare to equivalent stages in *C. elegans*. We gathered a total of 18 sample types, with 12 of those being region-specific and 6 being the whole worm. First, we validated our data by looking at the expression of well-known region-specific genes in *C. elegans* and then performed a comparative analysis within *C. elegans* young adult hermaphrodite to male^33^. We found that hermaphrodites were developing reproductive structures in comparison to males, which had increased metabolic and neuronal activity. The region-specific comparison demonstrated that we could detect differences in the worm in the absence of the gonads.

Next, we performed the first region-specific comparative analysis within *S. carpocapsae* young adults. We found that the female tail is enriched for neural development, and the head is regulating stimuli and sensing the environment. The enrichment in the tail region for neural activity in the first generation of *S. carpocapsae* female is a new finding, as neural activity in the tail is generally a focus in *C. elegans* males^107,108^. Within the Steinernematidae family, first generation females have unique tail projections, but there is no knowledge of neural activity in the tail, and if it is associated with reproduction or parasitism^109^. In contrast to females, *S. carpocapsae* males are developing structures for mating and also have heightened neural activity in the tail region. Lastly, we compared the young adults to IJs and found enrichment in neurogenesis in IJs, which might correspond to host seeking behavior and sensory responses^56^. The first comparative analysis within female and male young adults reveals transcriptomes of regions differ from one another and IJs, highlighting the importance of studying the profiles of all developmental stages.

Given all the data provided, we analyzed the transcriptomic differences between *S. carpocapsae* and *C. elegans*. The comparison between *S. carpocapsae* female to *C. elegans* hermaphrodites revealed genes enriched in *S. carpocapsae* female for energy regulation, fat storage, and several genes essential for vulva development. *S. carpocapsae* male compared to *C. elegans* male were enriched for metabolic activity but lacked neural related terms. The comparisons yielded shared GO terms such as “developmental process” which points to genes that may be at the core of developmental processes. These differences in core developmental processes, energy regulation and regulation of responses to stimuli may be the result of the evolutionary distance as well as lifestyle differences between the two species. While an entomopathogenic parasite such as *S. carpocapsae* is safely tucked away within a host during their development, adult life free-living species such as *C. elegans* face risks associated with changing environmental conditions and predators during their entire life cycle. This begs the question of whether these changes in expression patterns would vary similarly between parasites, EPNs, or other free-living nematodes as well as how these changes would fare in closely related nematodes such as other Steinernematids or Caenorhabditis clade members.

Studies of embryonic development between two Steinernematids including *S. carpocapsae* and two Caenorhabditis species including *C. elegans* have shown that embryonic development in *Steinernema* had notable differences in gene expression and delayed developmental timing and convergence at the late embryonic stage^57^. These differences in early development open speculation as to whether juvenile and early adult development is similarly delayed and how it maps to our findings of differences in gene expression at the young adult stage. Regardless, further investigation into conserved developmental genes necessary for the transition of young adults into the reproductive stage is of crucial significance for parasitism. Identifying core developmental genes would improve methods for biocontrol and mammalian parasite control. Lastly, we also compared IJ to dauer and found an increased GO enrichment of neurogenesis in dauers but not in IJs. This discrepancy raises questions about the conservation of neurons between two distantly related species.

This study provides the first transcriptional analysis of young adult and IJ with region-specific resolution in *S. carpocapsae* and the first comparison to the distantly related species *C. elegans*. We provided region-specific transcriptional profiles, a list of 115 genes essential for *S. carpocapsae* embryo and larval development, enrichment lists, differentially expressed gene analysis, GO analysis, and a gene-specific expression analysis of GO terms. Taken together, our results expand current knowledge of region-specific transcriptional profiles in *S. carpocapsae*. They offer an opportunity for the nematode community to use the data collected to advance studies in reproduction and life cycle, and link traits to transcriptome profiles, which subsequently leads to gene activity. Lastly, we believe these results are the first to instigate a more profound understanding of the conservation of nematode reproduction, neurogenesis, and development.

## Material and Methods

### Strains

*S. carpocapsae* (strain ALL) was cultured and maintained, according to Lu et al., 2017^16^. *C. elegans* Bristol strain N2 was used as the standard WT and strain him-8 were grown on Nematode Growth Media (NGM) plates seeded with OP50^110^.

### Experimental Design

We collected and sequenced single-nematode whole, heads, and tails for *S. carpocapsae* IJs, female and male adults. We also collected the equivalent for *C. elegans* dauers, males and hermaphrodites. We collected the mRNA and followed the steps in Serra et al., 2018. We sequenced a total of 54 samples with a sequencing depth average of approximately 10 million reads.

### *S. carpocapsae* IJs propagation

*S. carpocapsae* IJs were cultured and propagated *in vivo* using *Galleria mellonella* (waxworms) purchased from American Cricket Ranch (www.americancricketranch.com). Five wax worms were placed into a 10 cm petri dish with filter paper pressed to the bottom and 300 μL containing ~750 *S. carpocapsae* IJs (50 IJs/worm) was dispersed onto the filter paper. The infection plates were incubated at 25°C in the dark for 10 days. After the 10 days, the waxworm cadavers were transferred to White traps^111^. After 3 days, the IJs were collected, treated with Hyamine 1622 solution (Sigma cat# 51126) for 45 minutes and washed three times with Ringer’s. The IJs were stored at 25°C at a density of 10 IJs/ul in Ringer’s.

### Microscopy of *S. carpocapsae* female and male

*S. carpocapsae* males and females were washed from cultured plates and starved for 24 hours before imaging. Worms were anesthetized using 4μl levamisole and mounted on hydrocarbon oil. DIC images of females and males were collected on an upright Leica microscope with 5X and 20X. Images were stitched with Affinity Designer.

### *S. carpocapsae* culture and isolation of IJs, adult males and females whole, heads and tails

*S. carpocapsae* IJs (~10,000) were cultured with *Xenorhabdus nematophila* on a lipid agar (LA) plate (8 g/L of nutrient broth, 5 g/L of yeast extract, 2 g/L of MgCl2, 7 ml/L of corn syrup, 4 ml/L of corn oil, and 15 g/L of Bacto Agar) for up to 56 hours for adult females and males to avoid collecting fertilized females. Adults males and females were transferred from LA plates into a 1.5mL Eppendorf centrifuge tube. Adults were cleaned by washing twice with Ringer’s buffer (172 mM KCl, 68 mM NaCl, 5 mM NaHCO3, pH 6.1) and three times with Ultrapure DEPC water (Life Technologies cat# 750023. The pool of adults was then transferred to a slide that had been treated with RNAase away (VWR cat# 72830-022). IJs were washed as described above before proceeding to lysis. To collect whole adults and IJs, one clean adult/IJ was picked and transferred to a 0.2mL PCR tube and sliced in half with a 25-gauge needle^23^. To collect heads and tails from IJs and adults, a clean worm was transferred to a microscope slide (VWR cat# 16004-422) in 4μl of Ringer’s solution (made with DEPC water) + 0.01% Tween 20 and dissected under a stereomicroscope (Leica M80) with an X-ACTO knife (Amazon cat# B01MUWTH89). Head/tail was transferred to 0.2mL PCR tube with 2μL of lysis buffer, RNase inhibitor and proteinase K^23^.

### *C. elegans* culture and isolation of dauers, males, and hermaphrodites whole, heads, and tails

*C. elegans* were cultured with lawns of OP50 plates^110^. OP50 was cultured in Luria Broth (Thermo Fisher Scientific cat# 12780052) at 37 °C overnight and stored at 4 °C. 300 μl of OP50 were plated on NGM plates (17g/L Nutrient Agar, 3g/L NaCl, 2.5g/L peptone from animal tissue, 1mL/L 1M MgSO4, 1mL/L of 5mg/mL cholesterol in ethanol, 25mL/L 1M KPO4, 1mL/L 1M CaCl2)^112^. To collect RNA from whole adult males, heads and tails, *C. elegans* him-8(me4)IV plates were allowed to starve for 8 days at 25 °C, then chunked onto a NGM plate with lawns of OP50 and allowed to develop for 24 hours at 25 °C (*C. elegans* him-8 strain was provided by the CGC, which is funded by NIH Office of Research Infrastructure Programs (P40 OD010440)). Worms were transferred from the NGM plate into a 1.5mL Eppendorf centrifuge tube, then washed twice with M9 buffer and three times with Ultrapure DEPC water. The above steps were also used to collect whole hermaphrodites, heads, and tails but using *C. elegans* N2 (WT). To collect whole dauers, heads, and tails, *C. elegans* N2 (WT) were starved for sixteen weeks at 25°C. To confirm dauers phenotypically, nematodes were visualized on the EVOS inverted microscope for closed oral orifices by an internal plug, constricted pharynxes, and absence of pumping^112–114^. After washing nematodes, whole dauers and adults were transferred to a 0.2mL PCR tube and sliced in half with a 25-gauge needle according to Serra et al., 2018. To collect heads and tails from dauers and adults, a clean worm was transferred to a microscope slide (VWR cat# 16004-422) in 4μl of M9 (made with DEPC water) + 0.01% Tween 20 solution and dissected under a stereomicroscope (Leica M80) with an X-ACTO knife (Amazon cat# B01MUWTH89).

### RNA library preparation and sequencing

Samples collected followed the single-worm Smart-Seq2 protocol^23^. Briefly, samples collected in 2uL of lysis buffer were incubated at 65°C for 10 minutes and 85°C for 1 minute accordingly. Immediately following the incubation, 1μl of both dNTP and oligo-dT VN primers were added to the sample then incubated at 72°C for 3 minutes. Then, samples were reverse transcribed (RT) using a program set to 1) 42°C 90 min, 2) 50°C 2 min, 42°C 2 min (repeat 14x), 3) 70°C 15 min, 4) 4°C Hold. Then they were PCR amplified using a program set to 1) 98°C 3 min, 2) 98°C 20 sec, 67°C 15 sec, 72°C 6 min, (repeat 17x), 3) 72°C 5 min, 4) 4°C Hold. Amplified cDNA was cleaned with Ampure XP beads (Beckman Coulter cat# A63881) at a ratio of 1:1 (v/v). The concentration of samples was obtained with Qubit fluorometer, and cDNA quality checked in the Agilent 2100 Bioanalyzer. Samples that passed quality checks were tagmented with the Nextera DNA Library Prep Kit (Illumina cat# FC-121-1030). Tagmented samples were cleaned using the QIAquick DNA column (Qiagen cat# 28104). The tagmented cDNA was then amplified with barcodes using the Phusion High Fidelity PCR master mix (New England Biolabs cat# M0531L). The amplification program was set to 1) 72°C 5 min. 2) 98°C 30 sec. 3) 98°C 10 sec, 63°C 30 sec., 72°C 1 min. (repeat 10x). 4) 4°C Hold. The samples were cleaned with Ampure XP beads at 1:1 (v/v). Lastly, libraries were sequenced as paired-end, 43 base pair reads on the Illumina NextSeq 500.

### Gene expression Analyses

Genomes and GTF files were downloaded from WormBase Parasite (version WBPS14) for *S. carpocapsae* (PRJNA202318) and *C. elegans* (PRJNA13758). Both genomes were indexed with RSEM (version 1.2.25) using the command rsem-prepare-reference^115^. Unstranded, paired-end 43 bp RNA-seq reads were mapped using Bowtie^116^ (version 1.0.0) with the following options: bowtie -X 1500 -a -m 200 -S -p 8 -seedlen 25 - n 2 -v 3. After bowtie, gene expression was quantified with RSEM (version 1.2.25) with the following command: rsem-calculate-expression – bam – paired-end. For all analyses, gene expression was reported in Transcripts Per Million (TPM). We used counts for differential gene expression analysis.

The TPM file generated by RSEM for *S. carpocapsae* and *C. elegans* were normalized according to groups using the R package limma using command normalizeQuantiles^117^. Samples were normalized because they were collected, processed, and sequenced in different batches. Cluster 3.0 was used to generate heatmaps of gene expression with the following options: log transform, mean-centered genes, and hierarchically clustered^118^. The cdt file generated by cluster 3.0 were visualized with Java Treeview^119^. Heatmaps for differentially expressed genes were done with R package heatmap.2^120^.

Differential gene expression analysis was performed with edgeR and statmod^121,122^. First, RSEM count data were normalized by library size using calcNormFactors and then differential analysis was conducted with glmFit. Genes were considered differentially expressed if FDR < 0.05 and fold change > 2X. Differentially expressed (DE) genes found with edgeR and statmod were used to generate a TPM matrix. Volcano plots were generated with a list created from edgeR and TPM matrix with R packages ggplot2 and ggrepel.

### Orthology Relationship analysis

Protein sequences were downloaded from WormBase Parasite for *S. carpocapsae* and *C. elegans*. Then, protein sequences were filtered to keep the most extended isoform corresponding to each gene sequence. Orthology analysis was determined between *S. carpocapsae* and *C. elegans* using orthoMCL 1.4 with default settings^123^. For analysis, only one-to-one orthologs were used.

### Gene Ontology (GO) enrichment analysis

GO terms for *S. carpocapsae* were determined with Gene Ontology enrichment analyses using Blast2GO^124^ Fisher’s exact test and considered statistically significant if FDR < 0.05. Genes entered in Blast2GO were found DE by edgeR. For *C. elegans* analysis, GO terms were determined with Panther^125^ statistical overrepresentation test and considered statistically significant if FDR < 0.05. Genes entered into Panther were found to be DE by edgeR. Bubble plots were created using Revigo^126^ with default settings for *C. elegans*.

### Availability of Data

Data for this publication have been deposited in NCBI’s Gene Expression Omnibus (Edgar *et al*., 2002) and are accessible through GEO Series accession number GSE156356 (https://www.ncbi.nlm.nih.gov/geo/query/acc.cgi?acc=GSE156356).

## Acknowledgments

The authors thank the Initiative for Maximizing Student Development (IMSD) NIH Grant R25GM05524 and the Minority Access to Research Careers (MARC) Program, NIH Grant GM-69337.

---

**Supplemental Figure 1.**
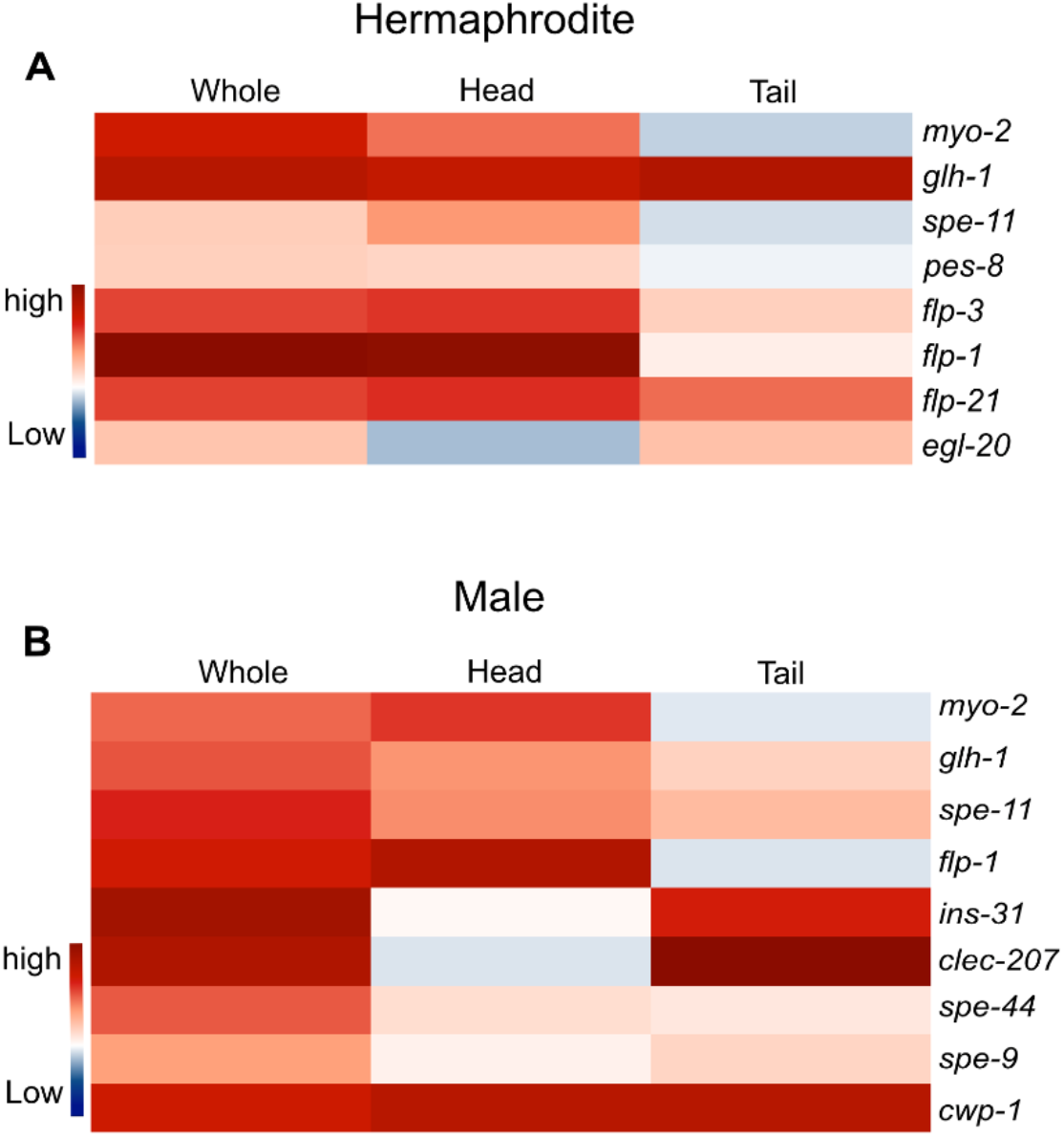
Validation of region-specific RNA expression in *C. elegans*. (a, b) RNA distribution correlates with regions collected for hermaphrodites and males.

**Supplemental Figure 2.**
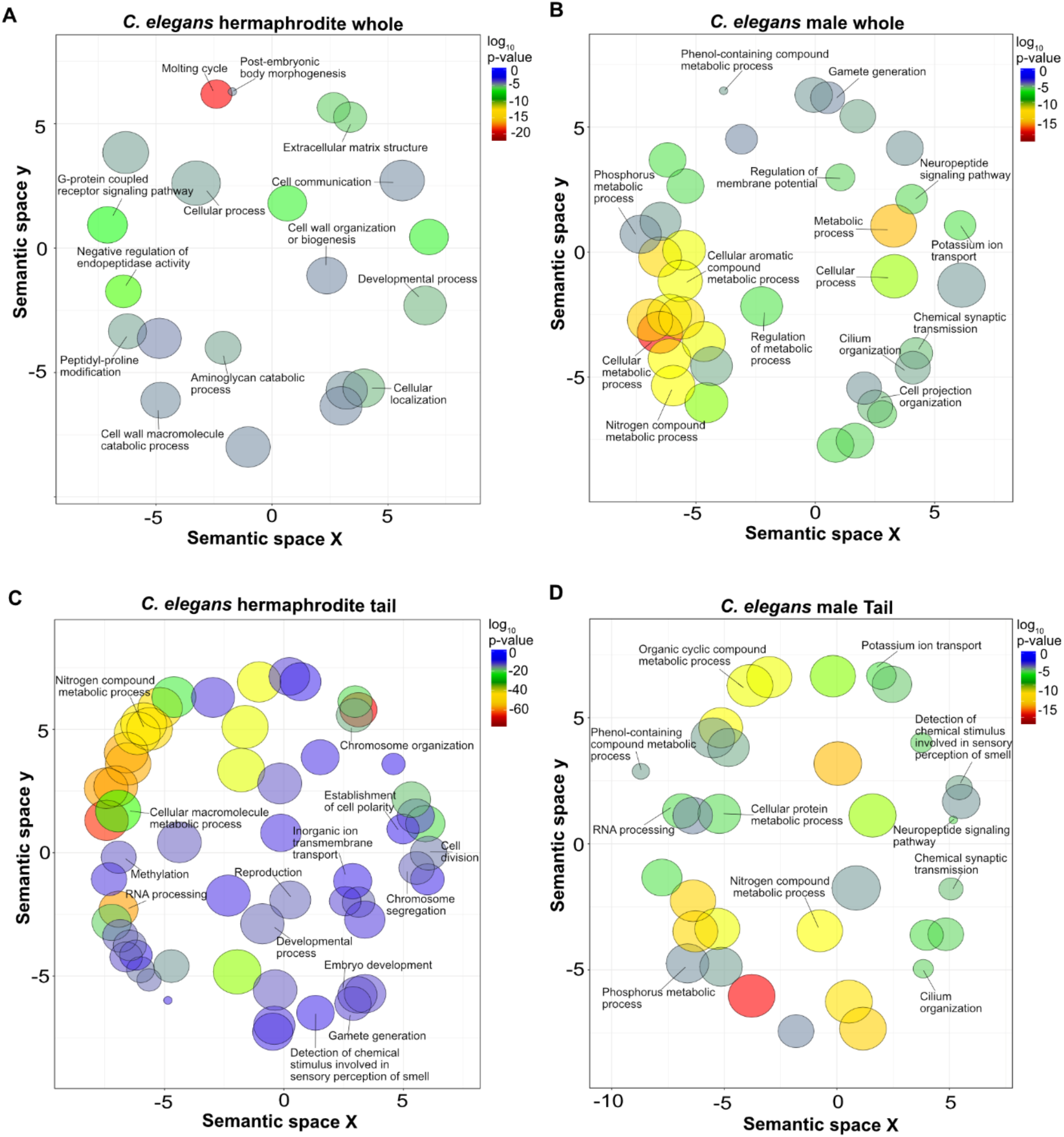
GO analysis of differentially expressed genes of *C. elegans* whole and tail of young adult hermaphrodite to whole and tail of young adult male. (a, b) DEGs from *C. elegans* young adult hermaphrodites and whole young adult male whole worms in Fig 2 panel B were subjected to biological processes GO analysis as in Fig 2E. (c, d) DEGs between *C. elegans* young adult hermaphrodite and young adult male tails from Fig 2 panel D were subjected to GO analysis as in Fig 2E.

**Supplemental Figure 3.**
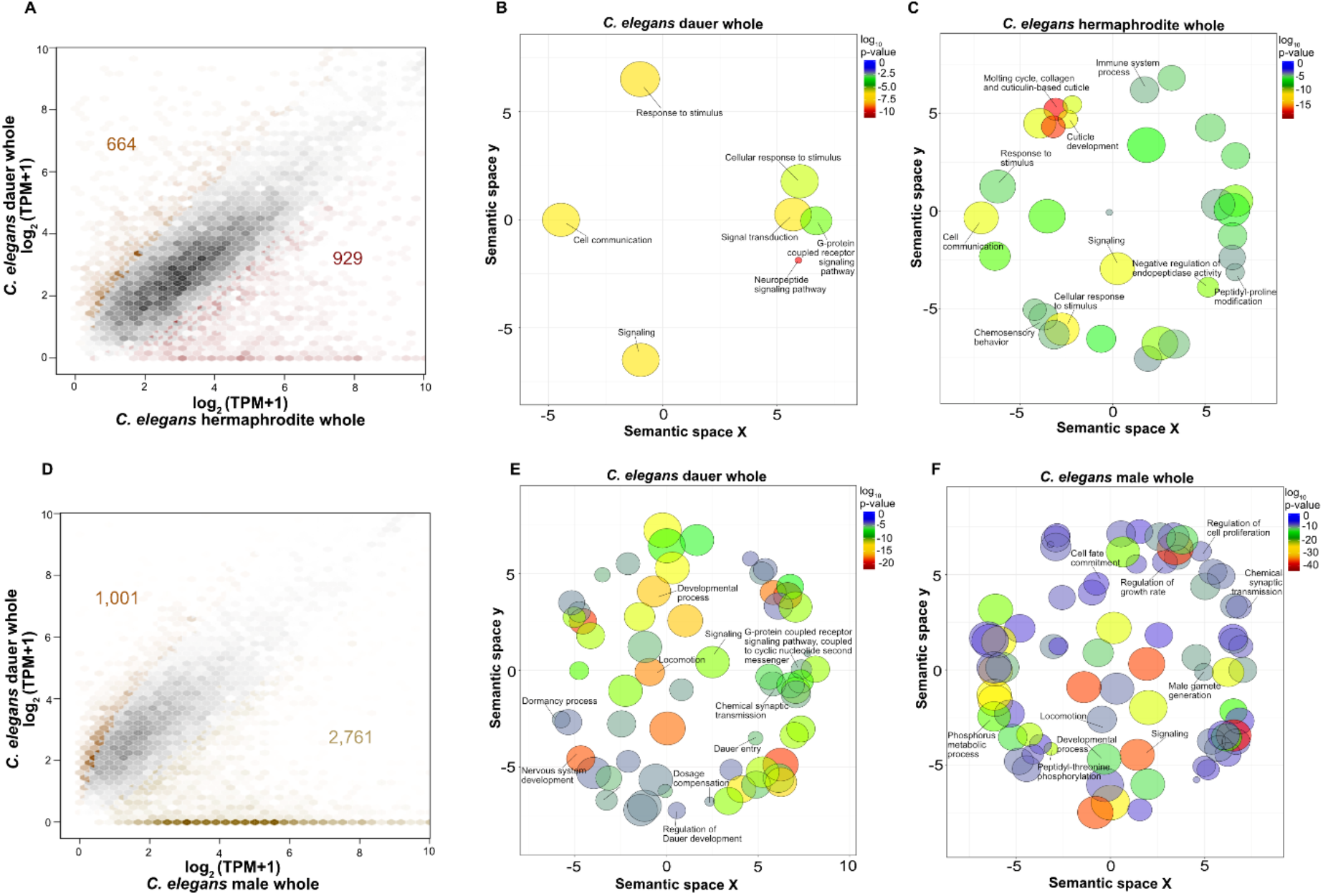
Analysis of differentially expressed genes in *C. elegans* whole dauers and young adult hermaphrodites and males. (a) Hexagonal-bins scatterplot of DEGs between *C. elegans* whole dauer and young adult hermaphrodites, depicting 664 genes DE in dauer compared to 929 in hermaphrodite. (b, c) DEGs from *C. elegans* whole dauers and whole young adult hermaphrodites in panel A were subjected to biological processes GO analysis (d) Hexagonal-bins scatterplot of the DEGs between *C. elegans* whole dauer and whole young adult male. (e, f) The DEGs between whole dauer and whole young adult males from panel D were subjected to GO analysis.

**Supplemental Figure 4.**
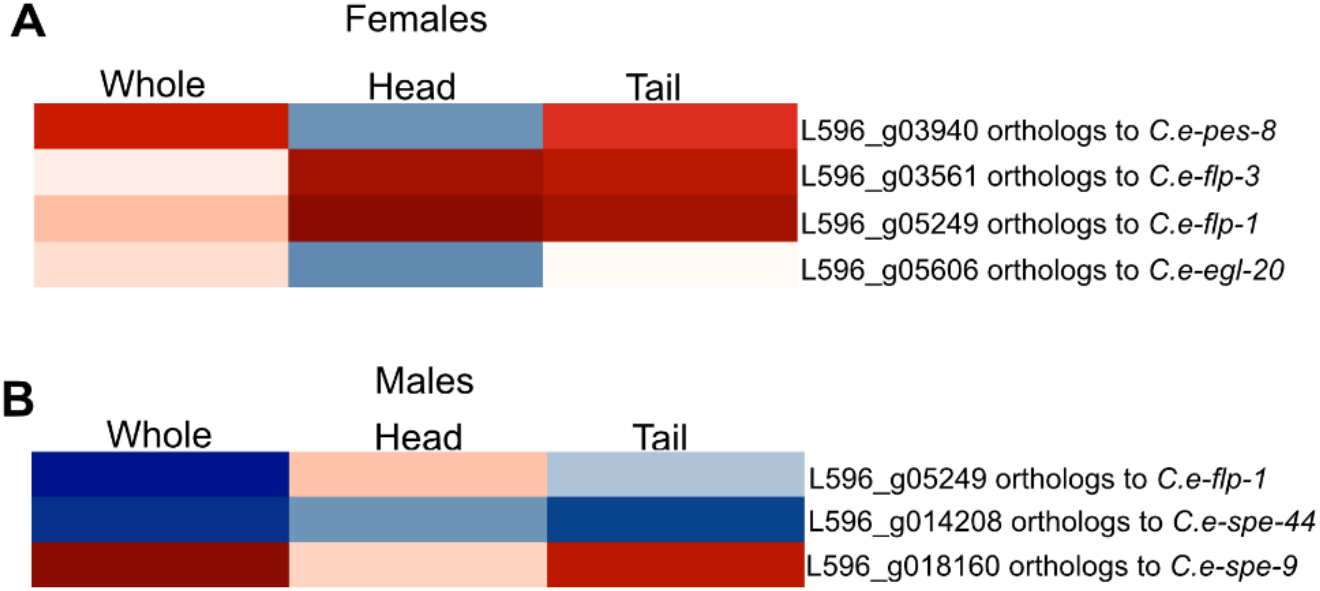
Expression of one-to-one orthologous genes of regionspecific RNA of *C. elegans* hermaphrodites and males observed in *S. carpocapsae* female and male. (a) RNA distribution of one-to-one of *C. elegans* hermaphrodite found in *S. carpocapsae* females. (b) RNA distribution of one-to-one of *C. elegans* male found in *S. carpocapsae* males.

**Supplemental Figure 5.**
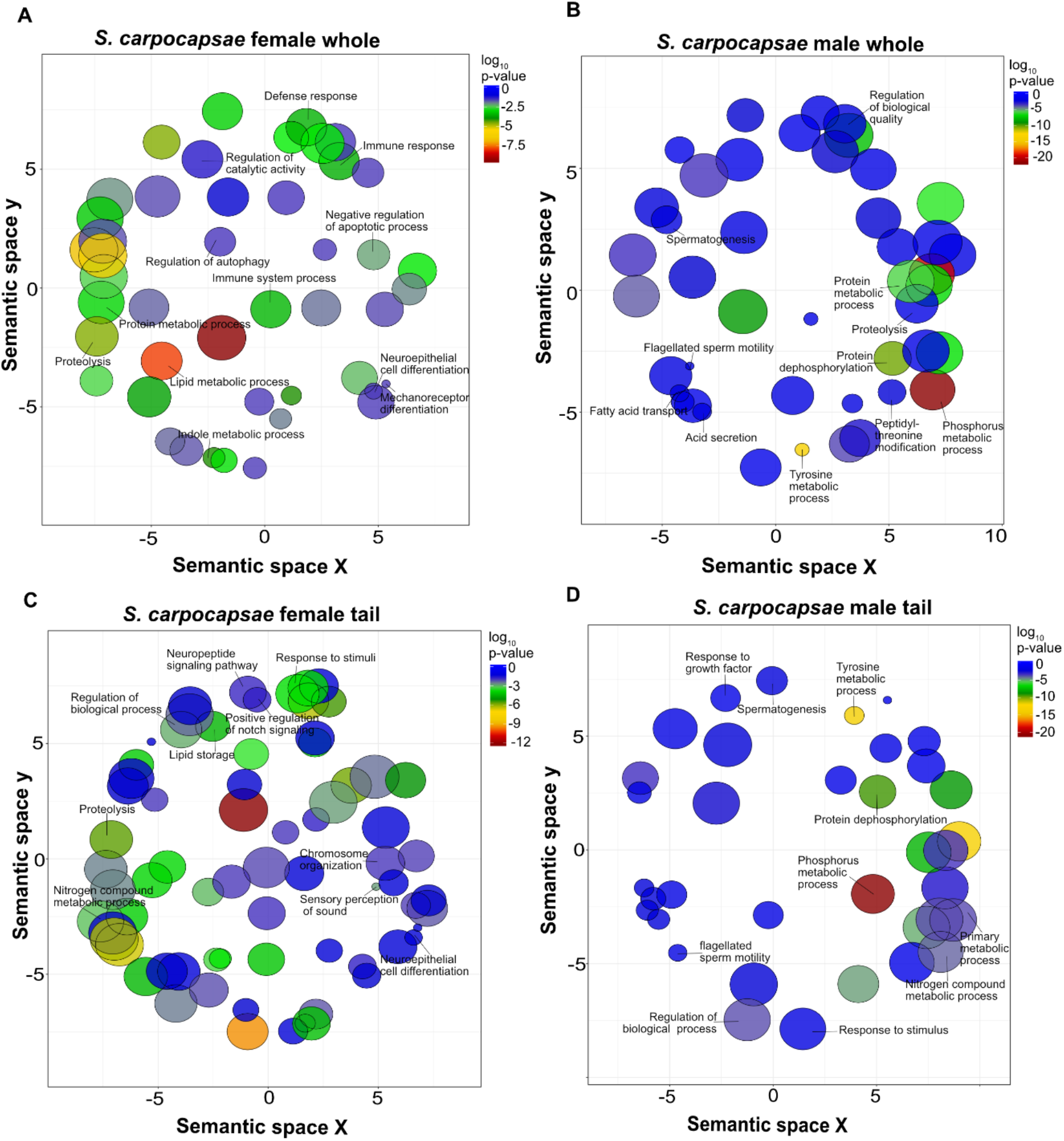
GO analysis of differentially expressed genes in *S. carpocapsae* young adult females and male whole nematode and tail. (a, b) GO analysis of DEGs from *S. carpocapsae* whole young adult female and males shown in Fig. 7B. (c, d) DEGs from *S. carpocapsae* young adult female and male tails depicted in Fig. 7D were subjected to GO analysis.

**Supplemental Figure 6.**
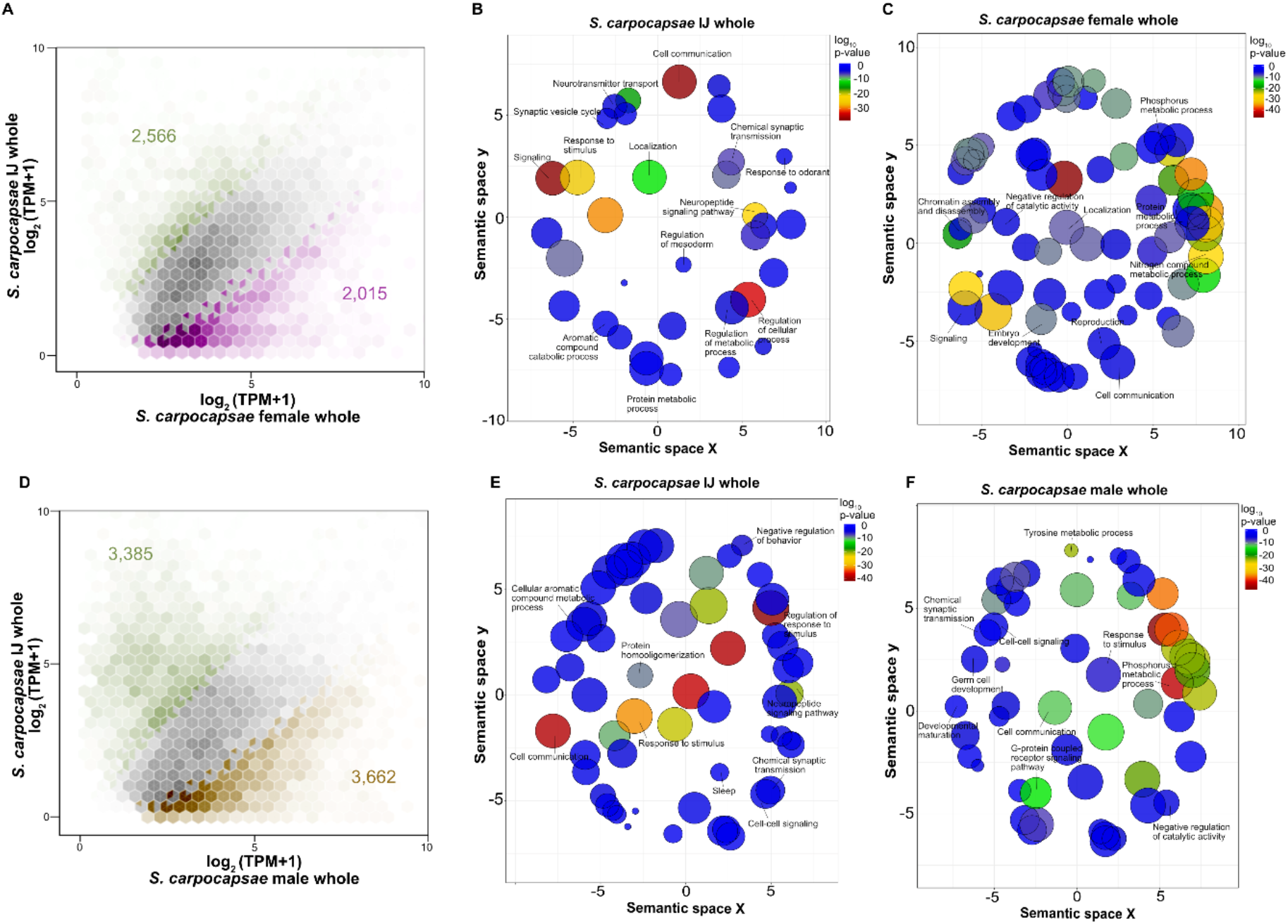
Analysis of differentially expressed genes in *S. carpocapsae* whole IJs to young adult females and males. (a) Hexagonal-bins scatterplot of DEGs between *S. carpocapsae* whole IJs versus young adult females. (b, c) DEGs from *S. carpocapsae* whole IJs and whole young adult female from panel A were subjected to biological processes GO analysis. (d) Hexagonal-bins scatterplot of the DEGs between *S. carpocapsae* whole IJs and whole young adult males. (e, f) The DEGs were between whole IJs and young adult males depicted in panel D were subjected to GO analysis.

**Supplemental Figure 7.**
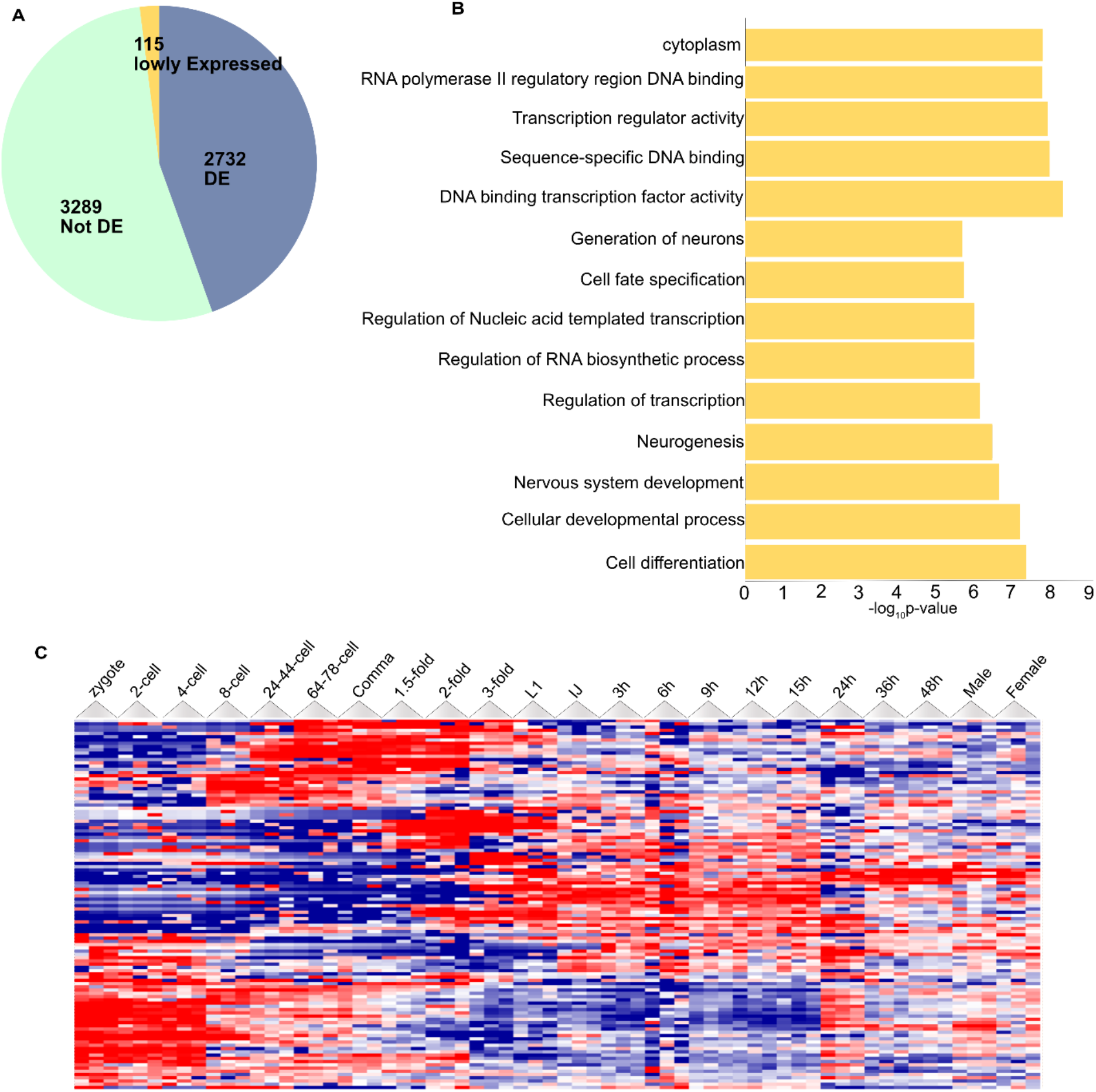
Gene expression analysis of one-to-one Orthologous genes not expressed at the young adult stage for *S. carpocapsae* and *C. elegans*. (a) Summary of all genes that are DE, not DE or not expressed in young adults. (b) GO analysis of the 115 lowly expressed genes. (c) Heatmap of the 115 genes through 20 developmental stages of *S. carpocapsae*.

**Supplemental Figure 8.**
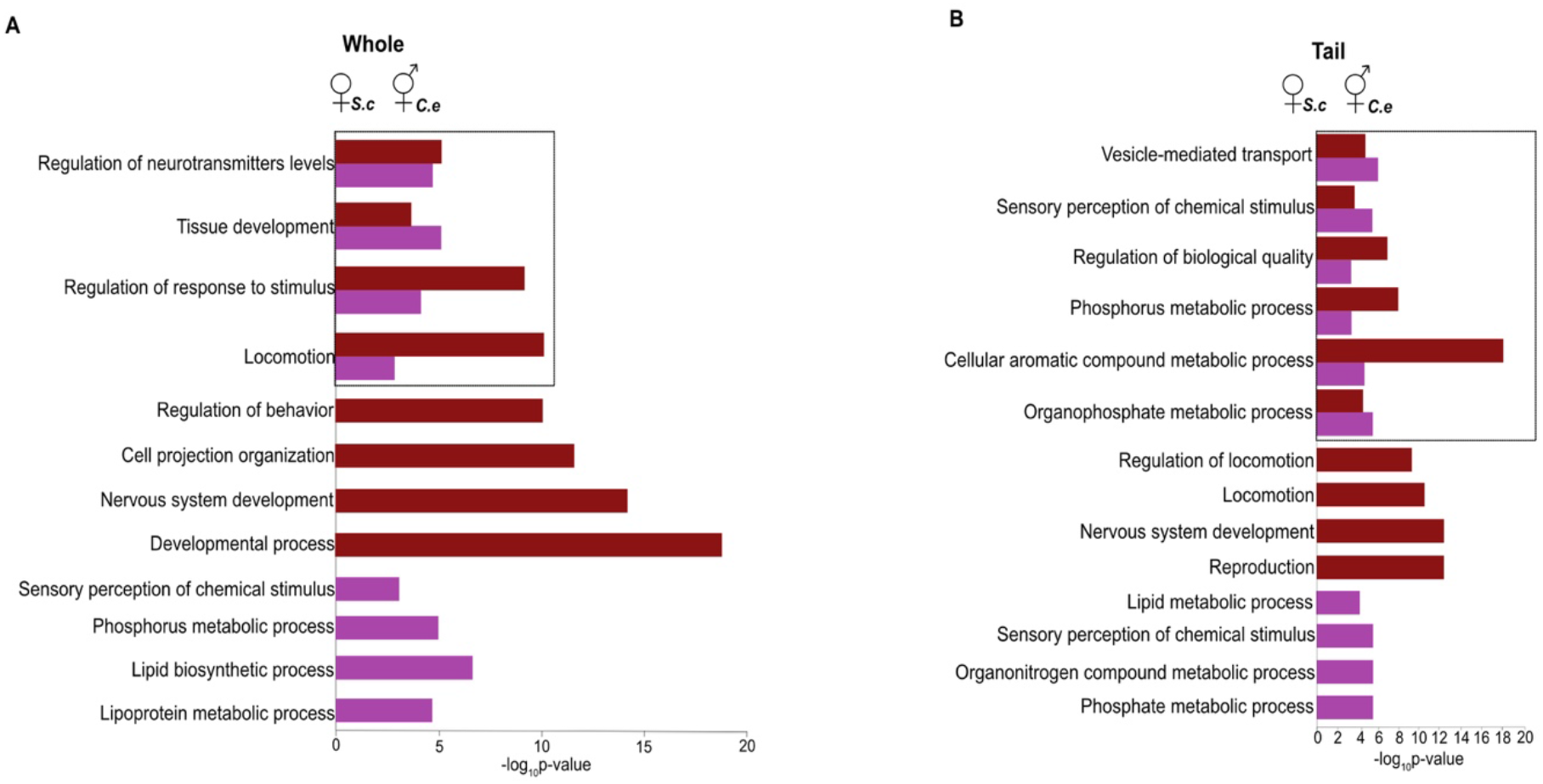
GO analysis of differentially expressed genes in *S. carpocapsae* young adult female and *C. elegans* young adult hermaphrodite whole nematode and tail. (a) Biological process GO terms for the DEGs between *S. carpocapsae* females (766 genes) and whole *C. elegans* hermaphrodites (1,097 genes) and (b) the DEGs between *S. carpocapsae* female (717 genes) and *C. elegans* hermaphrodite tails (635 genes). The bars are indicative of the P-value for the associated GO terms. Black box indicates shared GO terms.

**Supplemental Figure 9.**
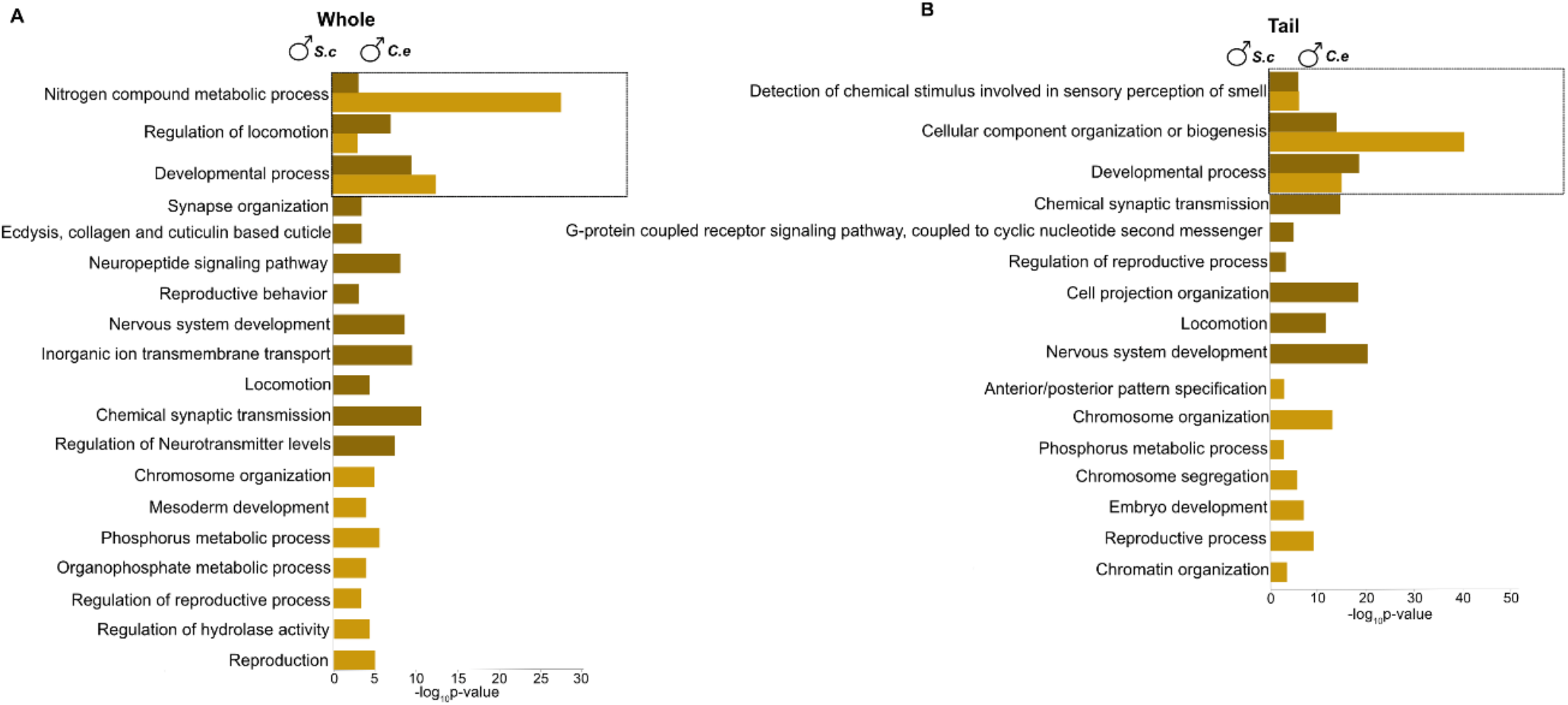
GO analysis of differentially expressed genes in *S. carpocapsae* young adult male and *C. elegans* young adult male whole nematode and tail. (a) Biological process GO terms for the DEGs between *S. carpocapsae* young adult male (919 genes) and *C. elegans* young adult male whole (1,047 genes) and (b) between *S. carpocapsae* male (1,109 genes) and *C. elegans* male tails (1,077 genes). The bars are indicative of the P-value for the associated GO terms. Black box indicates shared GO terms.

**Supplemental Figure 10.**
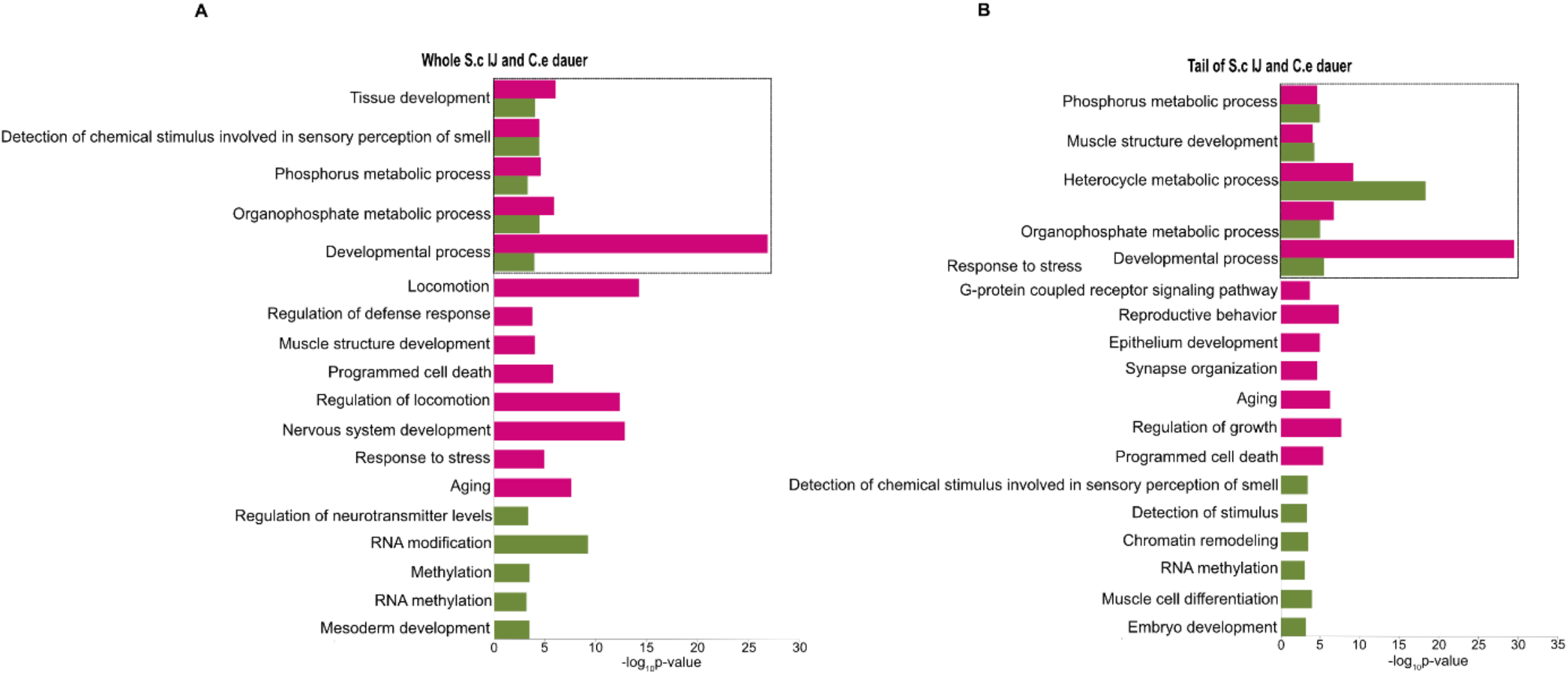
GO analysis of differentially expressed genes in *S. carpocapsae* IJ and *C. elegans* dauer whole nematode and tail. (a) The DEGs between *S. carpocapsae* IJ (1,289 genes) and *C. elegans* dauer (863 genes) and (b) *S. carpocapsae* IJ (1,011 genes) and *C. elegans* dauer tails (1,287 genes) were subjected to biological process GO analysis. The bars are indicative of the P-value for the associated GO terms. Black box indicates shared GO terms.

## Supplemental Tables

Table S1 *C. elegans* Young Adult Hermaphrodite vs Male Differentially Expressed Genes

Table S2 *C. elegans* Young Adult Hermaphrodite vs Male and Young Adult Hermaphrodite and Male vs Dauer Gene Ontology Analysis

Table S3 *C. elegans* Young Adult Hermaphrodite and Male vs Dauer Differentially Expressed Genes

Table S4 *S. carpocapsae* Young Adult Female vs Male Differentially Expressed genes

Table S5 *S. carpocapsae* Young Adult Female vs Male Adult and Young Adult female and Male vs Infective Juvenile Gene Ontology Analysis

Table S6 *S. carpocapsae* Young Adult Female and Male vs Infective Juvenile Differentially Expressed genes

Table S7 6136 1:1 orthologs between *S. carpocapsae* and *C. elegans*

Table S8 115 Developmental Ortholog genes not expressed in *C. elegans* and *S. carpocapsae* Young Adults

Table S9 Young adult *S. carpocapsae* female vs *C. elegans* hermaphrodite and *S. carpocapsae* male vs *C. elegans* male Differentially Expressed 1:1 ortholog genes

Table S10 Young adult *S. carpocapsae* female vs *C. elegans* hermaphrodite and *S. carpocapsae* male vs *C. elegans* male Gene Ontology Analysis

Table S11 *S. carpocapsae* Infective Juvenile vs *C. elegans* Dauer Differentially Expressed 1:1 ortholog genes

Table S12 *S. carpocapsae* Infective Juvenile vs *C. elegans* Dauer Gene Ontology Analysis

